# Information-theoretic Limits on Programmatic Specification of Biological Systems

**DOI:** 10.64898/2026.07.27.740886

**Authors:** Tuomo Kiiskinen, Oscar Kivinen, Manuel A Rivas

## Abstract

The central problem of biology is the origin of biological organization. We show, using information theory, that an organism does not contain enough organism-specific information to specify its own fully functioning microscopic organization. The organized machinery of life is therefore not the execution of a fully prewritten organism-specific program under favorable conditions. Rather, it is the compilation of a coarse organism-specific specification by a shared physical background that is constitutive of biological organization. We formalize this as a coarse-graining information threshold on biological specification, with two complementary entropy faces — a Shannon face controlling stochastic generation and a Hartley face controlling zero-error deterministic addressability. Above the threshold, organism-controlled information is sufficient to specify structural and functional ensembles; below it, programmed microstate determinism is impossible: deterministic addressability fails by pigeonhole, and any algorithm producing sub-threshold outputs must consume runtime randomness proportional to the information deficit. The threshold follows from two information-theoretic constraints — finite specification capacity and causal locality — supplemented by a mixing lemma showing that initial-condition information decays exponentially under thermal dynamics. We establish the threshold as a family of maximal capacity-compatible coarse-grainings, distinguish the proven impossibility below the threshold from the empirically realized coarse mappings above it, and locate the threshold empirically through worked cases of protein folding, *E. coli, Drosophila* early development, and *C. elegans*, together with computational verification using AlphaFold-2, the JCVI-syn3A 4D whole-cell simulation, and canonical stochastic gene network models. We further show that no known naturally realized environmental channel can close the gap. The result rules out *programmed microstate determinism* while leaving physical determinism untouched, reframes the genome as a generator specification rather than a trajectory program, and unifies gene-centric, developmental, and field-theoretic (bioelectric, morphogenetic, and related continuum) views of biological specification under a single coarse-graining framework.

## 1 Introduction

DNA was identified as the molecular substrate of heredity in the mid-twentieth century (Watson & Crick, 1953), and the genetic code — the mapping from nucleotide triplets to amino acids (Crick, 1970) — was solved shortly thereafter. At the level of protein primary structure, the encoding is now completely understood: given a coding sequence, one can read off the polypeptide it specifies with no ambiguity.

What is far less well understood is how the same molecule specifies *the rest* of the organism. A bacterium is not just a bag of proteins in stoichiometric ratios; it is a spatially organized, temporally coordinated, dynamically self-maintaining system. A metazoan is more striking still: a single fertilized cell gives rise to hundreds of distinct cell types arranged in a stereotyped body plan, with tissues, organs, and behaviors that recur reliably across individuals (Sulston *et al*., 1983). The mapping from sequence to amino acid is solved; the mapping from genome to organism is not.

A related information-scale argument has been made for neural wiring: the genome is far too small to specify a mammalian brain synapse by synapse, implying that innate architecture must be generated through compact developmental rules, architectural biases, and learning mechanisms rather than encoded as a wiring lookup table (Zador, 2019). Here we generalize this intuition from neural wiring to biological organization more broadly and formalize it as a coarse-graining threshold on organism-controlled specification.

Our core result is that the information budget available to a biological system — its genome plus continuous environmental signals — is fundamentally insufficient for the deterministic specification of its physical microstate trajectory. There exists a threshold coarse-graining *C*^∗^ such that organism-controlled information can specify the system at any resolution coarser than *C*^∗^, and cannot at any finer resolution. The threshold is fixed by the information capacity of the genome and environment relative to the entropy of the state space at each coarse-graining. Empirically, *C*^∗^ coincides with the resolution at which biological function is defined: copy numbers, expression states, and fate specifications lie above the threshold; spatial localization, conformational state, and exact temporal trajectories lie below it. The genome is a blueprint of the ensemble, not of the trajectory. Specification above the threshold is directly instantiated by mappings such as genome replication and the genetic code; the nontrivial question is where the finest reliably specified coarse-graining lies.

An analogy: the genome is not a figure-skating program choreographing every molecular movement, but a hockey coach’s game plan — roster, tactics, line assignments. The exact trajectory of the puck is computed by physics at runtime. The game looks like hockey because of the specified ensemble, not because every coordinate was pre-programmed.

We study *programmed microstate determinism* — whether the organism’s own specification scheme determines its trajectory at full resolution — and say nothing about *physical* determinism (whether the laws of physics determine the trajectory regardless of specification). The distinction is formalized in Definition 3.

## 2 Results

### 2.1 Formal physical framework

We formalize the organism as a physical information-processing system computing a microstate trajectory over an arbitrary temporal window [*t*_0_, *t*_*n*_] from a static genomic specification and a continuous stream of environmental inputs, analogous to an operating system computing a state trajectory from an executable and a stream of hardware interrupts. The system and its available information stores are defined through the following formal objects (Fig. 1):

**Figure 1:**
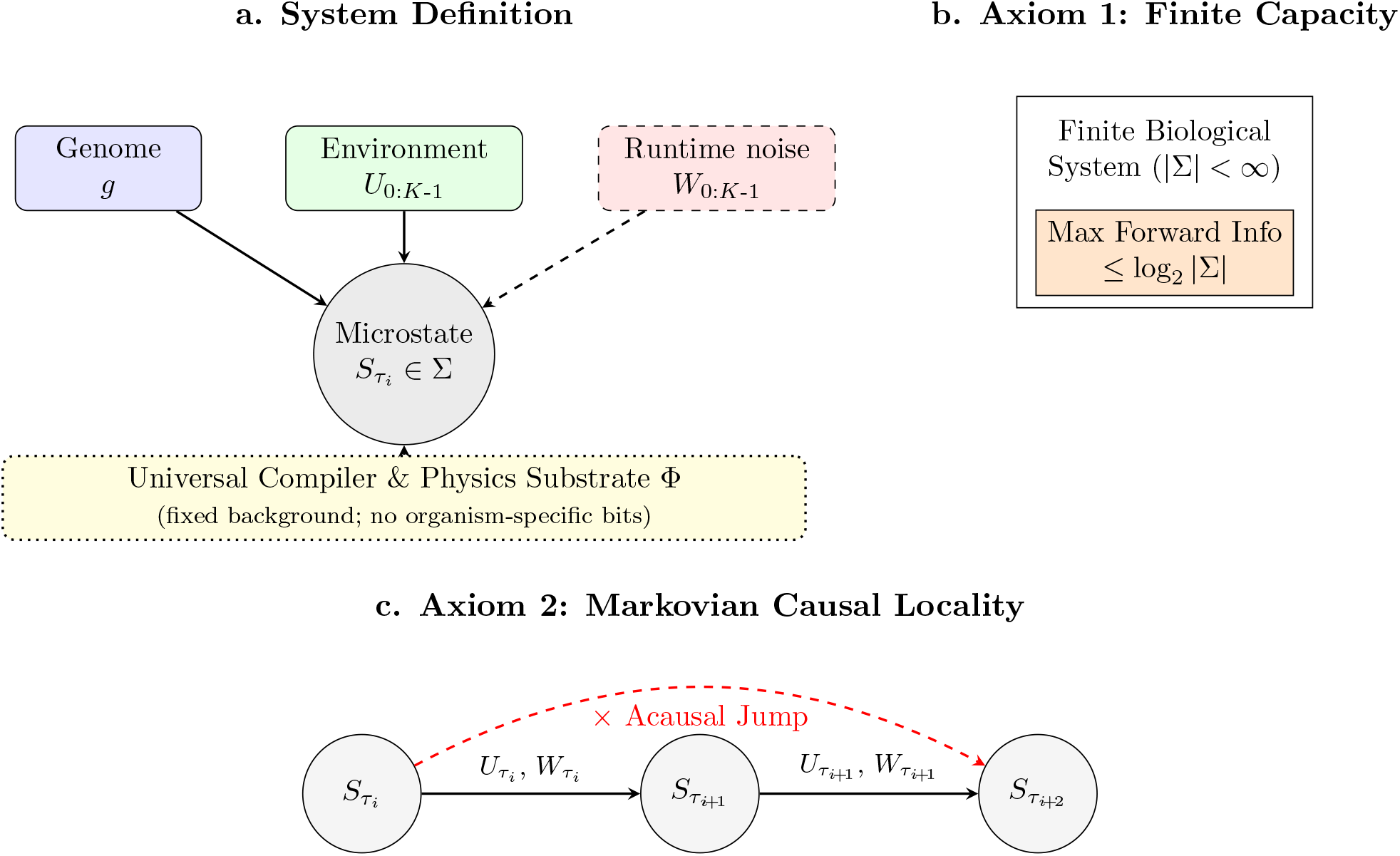
Formal physical framework of biological specification. **(a)** The physical microstate trajectory is computed from the static genome *g*, discretized environmental signals *U*_0:*K*−1_, thermal runtime randomness *W*_0:*K*−1_, and the universal compiler and physics substrate Φ. The genome, environmental channel, and runtime randomness are budgeted or runtime sources; Φ is held constant as fixed background (dotted border, no organism-specific bits) and is the medium through which the other three operate. **(b)** The physical constraint on forward state capacity. **(c)** Information can only propagate forward locally through intermediate physical states; acausal jumps are forbidden by the Markovian structure of physical dynamics.

#### 1. Microstate space (Σ) and physical trajectory

Let Σ be the physical state space at the resolution of full molecular microstate. At any instant, *S*_*t*_ ∈ Σ captures the exact spatial coordinates, conformational states, modification statuses, and binding partners of every molecule within the system boundaries. Σ is finite, being the Å-scale discretization of the compact configuration manifold of a bounded molecular system. Under a time-discretization *t*_0_ = *τ*_0_ *<* · · · *< τ*_*K*_ = *t*_*n*_ at resolution *δ*, the realized trajectory is 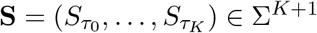. All entropies in this paper refer to finite random vectors of this form, computed under the probability space of Supplementary Note 1; impossibility results hold for every *δ >* 0.

#### 2. Genomic channel (*g*)

The static, onboard, heritable information store. A genome of *n* base pairs is a string over a four-letter alphabet; we define its specification capacity as the Hartley entropy *C*_*G*_ = log_2_ 4^*n*^ = 2*n*, i.e., the logarithm of the number of possible genome strings of that length. This is an upper bound on the Shannon entropy *H*(*g*) for any distribution over genomes, and it is the quantity computed in all empirical comparisons below. We use the full raw haploid sequence as a deliberately generous upper bound on organism-specific genomic capacity; functional capacity can only be smaller, so any threshold crossing against this budget is conservative and does not depend on classifying sequence as coding, regulatory, or inert. For the Hartley prong of the main theorem, *C*_*G*_ bounds the number of distinguishable messages the genome can carry; for the Shannon prong, it upper-bounds *H*(*g*) (see Supplementary Note 1).

#### 3. Continuous environmental channel 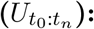

The integrated stream of exogenous physical inputs (chemical, mechanical, electromagnetic, thermal) crossing the system boundary during the interval. Under the time-discretization above, 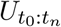 denotes the finite sequence 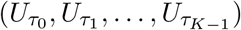, where *U* is the finite set of discretized environmental inputs at each step. Its integrated channel capacity is *C*_*E*_, bounded by the physical properties of the respective biological signaling modalities (Berg & Purcell, 1977). We retain the notation 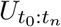 as a shorthand for the discretized sequence throughout.

#### 4. Runtime randomness (*W*_0:*K*−1_)

Ambient thermal fluctuations, quantum indeterminacy, and uncontrolled molecular collisions acting on the system through its coupling to the physics substrate. Under the time-discretization, 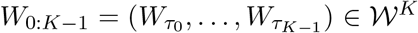 is a finite i.i.d. sequence. From the perspective of the organism’s specification scheme, it constitutes an unprogrammed source of bits not contained in 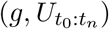.

#### 5. Universal compiler and physics substrate (Φ)

Two further objects are essential but are *not* signal channels and contribute no bits to the specification budget. The *universal compiler* is the invariant physical laws (electromagnetism, statistical mechanics, chemistry) that transform sparse specifications into organized microstates. The *physics substrate* is the specific physical regime under which those laws operate for a given biological system (solvent, temperature, thermal bath, pressure, ionic regime, hydrophobic effect). The laws are universal; the substrate is specific to life but invariant across individuals. Neither carries organism-specific information; neither is part of (*C*_*G*_ + *C*_*E*_). They are the precondition under which compilation proceeds, not contributors to the specification budget.

#### Continuous environmental channel and its capacity

Formally, the integrated channel capacity is 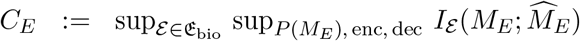, where E_bio_ is the class of biologically admissible boundary channels (each with source *M*_*E*_, encoder, receiver, and decoder 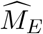) subject to rate, fidelity, addressability, synchronization, and biocompatible actuation constraints. If a modality admits an instantaneous rate envelope *r*_ℰ_ (*t*), then 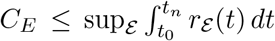 (the form used in Supplementary Note 2). The rate *r*_ℰ_ is addressed and decoded specifying information, not energy flux; broadcast fields contribute only a shared low-dimensional boundary condition unless they deliver distinct decodable messages to target cells.

The physical evolution of this system is governed by two undeniable structural axioms:

#### Axiom 1 (Finite state capacity)

Because Σ is finite, the maximum forward-control information that can be physically instantiated at any moment *t* is strictly bounded by log_2_ |Σ|.

#### Axiom 2 (Markovian causal locality)

The biological system obeys local physical dynamics. Under the discretization, 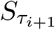 is conditionally dependent only on the immediately preceding state 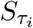, the environmental input 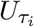, and the local runtime randomness 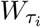. No forward-directed programmatic instruction can propagate from step *i* to step *j > i* + 1 without being physically instantiated in all intermediate states.

### 2.2 Formal definitions

We now formalize the threshold at which the specification budget is exhausted. The complete mathematical development, including the probability-space construction, technical assumptions, and proofs, is provided in Supplementary Note 1. Let *P* denote the lattice of all coarse-grainings of Σ.

#### Definition 1

(Coarse-graining). A coarse-graining *C* ∈ *P* of Σ is a partition of Σ into disjoint macroscopic cells. A coarse-graining *C*^*′*^ is finer than *C* (written *C*^*′*^ ≺ *C*) if every cell of *C*^*′*^ is contained in some cell of *C*.

#### Definition 2

(Induced trajectory and entropy). For a coarse-graining *C* of Σ, the induced discretized trajectory is 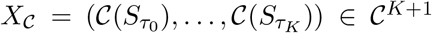. The entropy *H*(*X*_*C*_) is finite and is computed under the distribution induced by the physical dynamics (Supplementary Note 1). All coarse-grainings used in the empirical sections are pointwise partitions of Σ.

#### Remark 1

(Block-spin analogy and monotonicity). The projection **S** ↦ *X*_*C*_ is formally analogous to the block-spin (real-space renormalization group) operation of statistical physics: it is a many-toone map on discrete path space Σ^*K*+1^, and is therefore non-invertible. Passing from a finer coarsegraining *C*^*′*^ to a coarser one *C* (i.e., along *C*^*′*^ ≺ *C*) is well-defined and entropy-monotone, *H*(*X*_*C*_*′*) ≥ *H*(*X*_*C*_); passing from coarser to finer is not, since fine-grained paths are not recoverable from their coarse-grained projections. The lattice *P* of coarse-grainings inherits this one-way structure.

#### Proposition 1

(Monotonicity under refinement). *If C*^*′*^ ≺ *C, let*

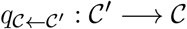

*be the surjection sending each fine cell to the unique coarse cell containing it. Then, denoting by π*_*C*←*C*_*′* : (*C*^*′*^)^*K*+1^ → *C*^*K*+1^ *the resulting map on trajectories*,

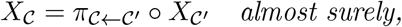

*and consequently*

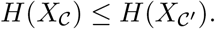

#### Definition 3

(Programmed microstate determinism). A biological system exhibits programmed microstate determinism at coarse-graining *C* if its realized trajectory *X*_*C*_ is determined by the joint specification 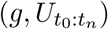. This admits two precise formulations:

- *Hartley form:* for every admissible specification (*g, u*), the conditional support is a singleton, | supp(*X*_*C*_ | *g* = *g, U* = *u*, Φ)| = 1. That is, the specification pins down a unique trajectory.
- *Shannon form:* the conditional Shannon entropy vanishes, 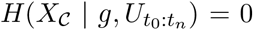, under the probability model of Supplementary Note 1. That is, *X*_*C*_ is almost surely a deterministic function of 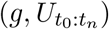.

The Hartley form implies the Shannon form. Both are properties of being specifiable by the organism’s own information stores. Both are distinct from physical determinism, which concerns whether the system’s microstate evolves deterministically under the laws of physics independent of any specification scheme. The theorem of this paper addresses programmed microstate determinism only.

### 2.3 Core lemmas

#### Lemma 1 (Runtime-randomness lower bound)

*Let A be any algorithm that, on input* 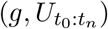, *reads i*.*i*.*d. fair random bits until halting and outputs X* ∈ Σ. *Let T denote the number of random bits consumed. Then for every fixed* 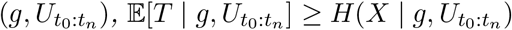.

This closes the algorithmic-compression loophole: a compressed program with a stochastic decompressor must consume runtime randomness at least equal to the output entropy. Bits can be relocated from explicit description into runtime randomness, but they cannot be made to vanish.

#### Connection to the physical dynamics

For a finite transition law, an exact prefix sampler can be constructed with expected fair-bit consumption less than 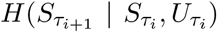 bits per step (Supplementary Note 1). Thus the expected runtime-bit consumption over *K* steps is controlled by the conditional path entropy; the thermal noise *W*_*t*_ is the physical source from which transitions draw bits not determined by (*g, U*).

Lemma 1 bounds randomness consumed given 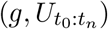. A deeper objection is that the full initial microstate 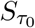 contains up to log_2_ |Σ| bits and could in principle seed a deterministic trajectory without consuming fresh randomness. Under a uniform KL contraction hypothesis for the discretized driven chain with contraction timescale 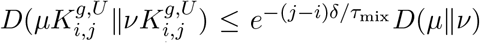 (a Dobrushin-type uniform ergodicity condition; Supplementary Note 1), this loophole closes:

##### Lemma 1b

**(Initial-condition mixing)**

*Under the uniform KL contraction assumption, the predictive mutual information carried by the initial microstate decays exponentially: for any step* 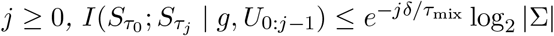.

#### Corollary (Runtime-randomness with initial-state side information)

*For any output X generated after j steps*, 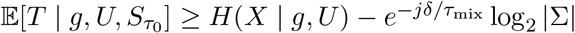.

For diffusive molecular degrees of freedom (Stokes-Einstein single-particle relaxation ∼ 10^−10^–10^−8^ s at body temperature, inflated to ∼ 10^−5^ s under the most adverse cytoplasmic-viscosity estimate), the exponent *jδ/τ*_mix_ exceeds 10^3^ on millisecond windows, so 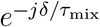 is astronomically small. The initial-condition loophole is closed for these fast residual modes.

##### Remark 2

(Scope of the mixing lemma). Lemma 1b closes the Laplacian-demon loophole for *fast* residual degrees of freedom. Long-lived structured memory (chromatin, epigenetic marks, slow condensates, membrane domains, cytoskeletal assemblies) is not claimed to mix on nanosecond scales; such slow variables are part of the current physical state itself and are governed by Lemma 2, not Lemma 1b. Slow biological memory is therefore not an initial-condition loophole; it is one of the finite stores whose capacity the framework bounds.

#### Scoped form of the mixing lemma

Formally, decompose 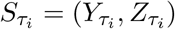 where *Y* is slow state-carried memory and *Z* is the fast residual. Under uniform KL contraction *only* for the conditional fast dynamics, 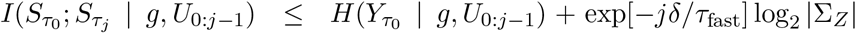. The first term is slow state-carried memory governed by causal locality (Lemma 2); the second is the fast initial-condition residue, which decays.

##### Lemma 2

**(Causal locality)**

*Let* 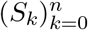 *be the discretized trajectory on* Σ, *Markov in S*_*k*_ *conditional on* (*S*_*k*−1_, *U*_*k*_, *W*_*k*_). *For any measurable forward program A*_*k*:*n*_ = *ϕ*(*S*_*k*−1_, *U*_*k*:*n*_) *and current-function readout f* (*S*_*k*−1_), *H*(*f* (*S*_*k*−1_), *ϕ*(*S*_*k*−1_, *U*_*k*:*n*_) | *U*_*k*:*n*_) ≤ *H*(*S*_*k*−1_ | *U*_*k*:*n*_) ≤ log_2_ |Σ|.

The lemma defines “programmed”: a forward action is programmed iff it is a deterministic function of the current state and future environmental inputs, with no dependence on future thermal noise. The lemma is compatible with biological state-carried memory (phosphorylation, epigenetic marks, synaptic weights, membrane potential are all part of *S*_*t*_ and counted by the same bound); what it forbids is delayed instructions indexed to wall-clock time without a state-borne trigger. DNA is compatible as a *generator specification* but not as a free delayed-trajectory tape, since transcription is initiated by local promoter recognition, polymerase availability, and chromatin accessibility (Munsky *et al*., 2012) — all state variables — rather than by a global clock indexing the genome. Supplementary Note 1 records the conditional-independence structure.

### 2.4 Zero-error addressability and the cost of reducing a fine-state fiber

A third constraint, complementary to the runtime-randomness and causal-locality lemmas, governs the *deterministic addressability* of fine states: how many bits an organism-controlled specification must expend to select an arbitrary fine-grained realization within the fiber left unresolved by a functional coarse-graining. This is a zero-error coding statement, distinct from Shannon entropy, and is the appropriate object whenever the empirical claim concerns deterministic specification rather than stochastic sampling.

#### Definition 4

(Admissible fiber and zero-error support cost). Fix the organism-controlled specification together with the shared physical background, 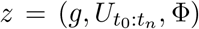. For a coarse-graining *C*, define the admissible fine-state fiber *A*_*C*_(*z*) = supp (*X*_*C*_ | *z*), and the zero-error support cost *H*_0_ (*X*_*C*_ | *z*) = log_2_ | *A*_*C*_(*z*)|. This is the bits required to address an arbitrary distinguishable finegrained realization in the fiber, distinct from *H*(*X*_*C*_ | *z*), with *H* ≤ *H*_0_ in general and equality iff the conditional law is uniform.

#### Proposition 2

(Zero-error addressability bound). *Fix* 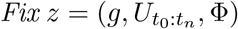, *If a deterministic specifier uses a message M of length at most L bits and side information R to select states in A*_*C*_(*z*) *with zero error, then L* + *H*_0_ (*R* | *z*) ≥ *H*_0_ (*X*_*C*_ | *z*) .

#### Corollary 1

(Cost of reducing a fine-state fiber). *If a state-carried variable R*_*t*_ ⊆ *S*_*t*_ *reduces* max_*r*_ log_2_ |*A*_*C*_(*z, r*)| ≤ *h, then H*_0_(*R*_*t*_ | *z*) ≥ *H*_0_(*X*_*C*_ | *z*) − *h. Any biological mechanism that reduces the unresolved fine-state support by β bits must therefore physically carry those β bits in g, U, or the current state S*_*t*_, *where they are counted by the specification budget and by causal locality*.

The corollary identifies precisely where any putative “fiber-reducing” mechanism must reside: in *g*, in *U*, in the current state *S*_*t*_ (subject to Lemma 2), in the universal compiler Φ (shared, not organism-specific), or in runtime randomness (not programmed). There is no fourth category. Combinatorial calculations in the empirical sections below should be read as zero-error addressability bounds on *H*_0_(*X*_*C*_ | *z*), not as lower bounds on Shannon entropy.

### 2.5 The coarse-graining threshold

#### Notation: budgeted message and fixed background

Separate the organism-controlled budgeted message 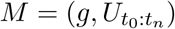 from the fixed, non-message background *b* = (Φ, system identity, [*t*_0_, *t*_*n*_], *δ*), which contributes no organism-specific bits. The total specification budget is *B* = *C*_*G*_ + *C*_*E*_, with | supp(*M*)| ≤ 2^*B*^ for the Hartley prong and *H*(*M*) ≤ log_2_ | supp(*M*)| ≤ *B* for the Shannon prong (Supplementary Note 1).

#### The unified threshold

For a generic information cost ℋ ∈ {*H, H*_0_}, the capacity-compatible coarse-grainings are C_ℋ_(*B*) = *{C ∈ P* : *ℋ*(*X*_*C*_ | *b*) ≤ *B*}, with antichain 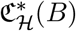 of maximal elements under refinement. The two specializations are the *Shannon* face C_1_ (with ℋ= *H*, governing runtime randomness for stochastic generation) and the *Hartley* face C_0_ (with ℋ= *H*_0_, governing zero-error deterministic addressability). Since *H* ≤ *H*_0_, C_0_(*B*) ⊆ C_1_(*B*); zero-error compatibility implies Shannon compatibility, and the zero-error threshold is at least as coarse. Between the two lies a biologically meaningful zone where stochastic ensemble generation fits the message budget but zero-error addressability does not.

##### Theorem 1

**(Coarse-graining threshold for programmed specification)**

*Let P be the lattice of coarse-grainings of* Σ *ordered by refinement, let b denote the fixed non-message background, let* 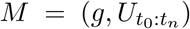 *denote the budgeted organism-controlled message with* | supp(*M*)| ≤ 2 *where B* = *C*_*G*_ + *C*_*E*_. *Then:*

1. *(Capacity-compatible regime.) If C ∈* C_1_(*B*), *then stochastic specification at resolution C is not ruled out by Shannon capacity alone: there exists an abstract lossless source code for X*_*C*_ *whose expected code length is at most H*(*X*_*C*_ | *b*)+1 ≤ *B* +1 *bits. If C ∈* C_0_(*B*), *then zero-error deterministic addressability is not ruled out by Hartley capacity alone. The inequalities alone do not establish realization; sufficiency for a particular coarse-graining is established when the corresponding code is present in M, causally exposed through state, and physically realized — as in genome replication and the genetic code*.
2. *(Sub-threshold impossibility, two prongs*.*) Let C*^*′*^ *be a strict refinement below the relevant threshold antichain*.
  a. *Zero-error addressability: if C*^*′*^ *is a strict refinement of some* 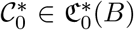 *and H*_0_(*X*_*C*_*′* | *b*) *> B, then no organism-controlled deterministic specification of capacity B can address every distinguishable fine state in* supp(*X*_*C*_*′* | *b*).
  b. *Runtime-randomness cost: if C*^*′*^ *is a strict refinement of some* 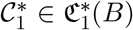 *and H*(*X*_*C*_*′* | *b*) *> B, then H*(*X*_*C*_*′* | *M, b*) ≥ *H*(*X*_*C*_*′* | *b*) − *B, and any algorithm producing X*_*C*_*′ from M must consume runtime randomness with expected length at least H*(*X*_*C*_*′* | *b*) − *B*.

##### Remark 3

(Approximate addressability). Both faces relax under tolerated error probability *ε*: by Fano’s inequality, the bound softens by *O*(*h*_2_(*ε*)+*ε* log *K*). For biological gaps that exceed organism-controlled capacity by factors of 10 or more, the order-of-magnitude impossibility is intact.

##### Remark 4

(What a sufficiency proof would require). Capacity compatibility becomes sufficiency through a *realizable causal code*: a message *M* with *H*(*X*_*C*_ | *M, b*) = 0 that is causally exposed through state and physically decoded by the Markov dynamics. Such codes are empirically realized for coarse mappings including genome replication and the genetic code. For arbitrary trajectory-level coarse-grainings no general construction is claimed; the impossibility results below the threshold do not depend on such a construction.

The threshold is in general not a single canonical coarse-graining but an antichain of maximal compatible coarse-grainings; the empirical claim is that biological function is defined at coarse-grainings lying within these antichains.

#### Notation for empirical comparisons

Throughout the empirical sections we use the raw haploid genome bit count 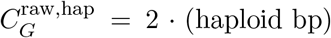 as a generous upper bound on *C*_*G*_. For diploid organisms this is also a tight upper bound on the diploid information content, to within the withinindividual heterozygosity rate 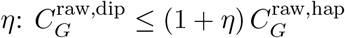 by the chain rule of entropy, with *η <* 0.01 in humans (1000 Genomes Project Consortium, 2015) (full derivation in Methods). Functional capacity satisfies 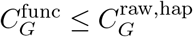, so using 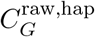 is conservative. Even the narrowest threshold crossing reported below exceeds the plausible ploidy correction by at least an order of magnitude, and the larger coding-input and trajectory gaps are wider still. In tables that mix Shannon and Hartley entropies, descriptor type is specified per row.

#### Corollary (Ensemble specification is forced)

*Below the threshold, biological function must be realized as a property of the conditional distribution P* (*X*_*C*_*′* | *M, b*) *rather than as a property of individual realizations specified by the organism*.

### 2.6 Protein folding

The genetic code is a deterministic 1:1 mapping from nucleotide triplets to amino acids, so the genome exactly specifies the primary sequence of every protein the organism produces. Primary sequence, however, is a one-dimensional object: a string over a twenty-letter alphabet. The biologically functional object is the three-dimensional folded structure, and the information required to specify a 3D structure is vastly greater than the information required to specify its sequence.

A protein of ∼ 300 residues carries ∼ 300 log_2_ 20 ≈ 1300 sequence bits, equivalently ∼ 1800 nucleotide bits at three nucleotides per residue. A conservative backbone-level 3D specification at ∼ 1 Å precision requires ∼20 bits per residue (three coordinates at ∼7 bits each), giving ∼ 6 × 10^3^ bits per protein. Atomic-resolution specification including side chains (∼8 heavy atoms per residue) raises the per-protein cost to ∼170 bits per residue, or ∼ 5 × 10^4^ bits. These static estimates remain conservative because a functional protein occupies a conformational ensemble; specifying its accessible structural states and characteristic motions requires additional information beyond any single coordinate snapshot.

Even the protein backbone alone exceeds the information carried by its coding sequence once specified beyond a very coarse resolution. Consider an aggressively compressed internal-coordinate description that fixes bond lengths, bond angles, and peptide-bond geometry, removes global translation and rotation, omits all side chains, and retains only the two backbone torsions *ϕ* and *ψ* per residue. At angular resolution Δ*θ*, its direct structural cost is

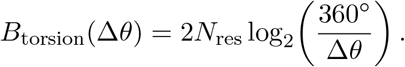

The coding sequence contributes six raw nucleotide bits per residue, so equality occurs only at Δ*θ* = 45^°^. Any finer backbone specification therefore exceeds the exonic nucleotide information, even before side-chain geometry, deviations from ideal bond geometry, alternative conformations, modifications, or dynamics are included.

Aggregated across the *E. coli* proteome (∼4288 protein species), the total atomic-resolution specification cost is 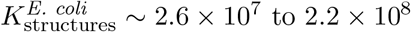 bits, exceeding the raw haploid genomic budget of ∼ 9.3 × 10^6^ bits by a factor of 3 to 24. For the human proteome (23,586 reviewed protein species, ∼ 1.5 × 10^7^ residues), the aggregate at atomic resolution is ∼ 2.5 × 10^9^ bits, ∼41% of the raw human haploid genome (∼ 6.1 × 10^9^ bits); at backbone resolution the aggregate is ∼ 3 × 10^8^ bits, ∼5% of the genome.

The threshold *C*^∗^ for a protein therefore sits between the primary sequence (∼ 10^3^ bits, above; genomically specified) and atomic-resolution coordinates (∼ 10^4^–10^5^ bits per protein, below; aggregate cost exceeds the genome). Folded structure is not stored as organism-specific coordinate information; the genome supplies the sequence, and the physical substrate compiles the structural ensemble. A natural objection is that folded proteins, once compiled, reduce later cellular spatial-organization fibers through binding interfaces and scaffolds; but by the zero-error addressability bound, any reduction of *q* bits requires *q* bits of side information physically instantiated in the current state, competing with current function for the same finite log_2_ |Σ| budget. Compiler-mediated reduction is not an uncounted genomic blueprint.

### 2.7 Information thresholds in model organisms

#### E. coli

The reference genome (4,639,221 bp) gives *C*_*G*_ ≤ 9.28 × 10^6^ bits. For *S* ≈ 4288 protein species with total count *M* ≈ 4 × 10^6^ (Taniguchi *et al*., 2010; Milo *et al*., 2010), the copy-number vector has 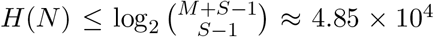 bits, two orders of magnitude below the genomic budget. A conservative 4 nm-occupancy spatial descriptor for the proteome gives *H*_0_ ≈ 1.2–1.5×10^7^ bits, exceeding the budget. By Theorem 1 part 2(a), no deterministic specification at capacity *C*_*G*_ can address every distinguishable spatial configuration.

#### Drosophila

For the best-measured biological environmental channel, the Bicoid morphogen carries ∼1.5 bits per nucleus per fresh readout, with recovery time *τ*_*c*_ ≈ 69 s (Gregor *et al*., 2007; Tkačik *et al*., 2008; Petkova *et al*., 2019). Over a one-hour window (∼53 independent samples), the per-nucleus environmental budget is ∼80–230 bits — sufficient to specify gap-gene fates but not a developmental microstate trajectory. The environmental channel is itself an ensemble specifier.

#### C. elegans

The genome (∼100.3 Mb, *C*_*G*_ ≤ 2.006×10^8^ bits) easily accommodates static and coarse dynamic descriptors of development but is exceeded by exact morphological microstates. Across a developmental descriptor ladder (Table 1), static symbolic descriptors (lineage/fate/timing, 8.64 × 10^3^ bits; adult connectome with 8 bits/edge, 4.37 × 10^4^ bits) and coarse dynamic descriptors (a nuclear-position movie over 558 nuclei at 0.1 *µ*m resolution sampled minutely over 16 h, 1.31 × 10^7^ bits, 6.5% of *C*_*G*_; a contact-graph movie, 9.81 × 10^5^ bits) all remain capacity-compatible. The threshold appears only when descriptors are refined toward exact dynamic morphology: a scalar morphology movie reaches *C*_*G*_ at ∼1530 bits/cell/frame, voxel cell-boundary addressability crosses between *δ* = 0.25 *µ*m and 0.10 *µ*m, and a full egg-to-adult dynamic trajectory crosses at 41.5–71.3 bits/cell/min (959 adult somatic cells over 5040 min). The embryogenesis-only crossing is higher, ∼374 bits/cell/min over 960 min for 558 nuclei. Empirically, ∼43% of cell–cell connections vary across isogenic individuals (Witvliet *et al*., 2021), confirming that the connectome is not specified to single-trajectory precision (Zador, 2019).

**Table 1:** *C. elegans* descriptor ladder. Costs are zero-error addressability quantities *H*_0_, except threshold rows, which report the per-unit cost at which the descriptor equals *C*_*G*_ = 2.006 × 10^8^ bits.

| Descriptor | Bits or threshold | Ratio to $C_G$ |
| --- | --- | --- |
| Symbolic lineage/fate/timing | $8.64 \times 10^3$ | $4.3 \times 10^{-5}$ |
| Static connectome, 8 bits/edge | $4.37 \times 10^4$ | $2.2 \times 10^{-4}$ |
| Nuclear-position movie | $1.31 \times 10^7$ | $6.5 \times 10^{-2}$ |
| Contact-graph movie | $9.81 \times 10^5$ | $4.9 \times 10^{-3}$ |
| Scalar morphology, 100 bits/cell/frame | $1.31 \times 10^7$ | $6.5 \times 10^{-2}$ |
| Scalar morphology crossing | 1530 bits/cell/frame | 1.0 |
| Voxel segmentation, $\delta = 0.10 \mu\text{m}$ | $2.20 \times 10^8$ | 1.10 |
| Egg-to-adult trajectory crossing | 41.5–71.3 bits/cell/min | 1.0 |
| Embryogenesis-only trajectory crossing | 374 bits/cell/min | 1.0 |

In each organism, functional and coarse-dynamic descriptors lie at least two orders of magnitude below the genomic budget, while exact microstate or exact dynamic trajectory descriptors exceed it.

### 2.8 Physical impossibility of the continuous environmental rescue

The addressability deficit at a sub-threshold coarse-graining, Δ_0_ = [*H*_0_(*X*_*C*_*′* | *b*) − (*C*_*G*_ + *C*_*E*_)]_+_, fixes the bit rate any rescue channel would need to deliver. For *E. coli*, the representative deficit of ∼ 5 × 10^6^ bits over a 30-minute generation requires approximately 3 × 10^3^ addressed bits s^−1^ per cell. Under the bacterial parameters of Supplementary Note 2, and using assumptions deliberately generous to the rescue hypothesis — that is, chosen to maximize the information credited to the environmental channel — an idealized Berg–Purcell/Gaussian-equivalent calculation gives approximately 40 bits s^−1^ per independent scalar ligand channel. More generally, no known naturally realized source–receiver architecture examined here simultaneously provides the required rate, microstate addressability, feedback, synchronization, and actuation depth. Engineered external controllers are not ruled out; they enlarge the specification system and must pay the missing information cost through an enlarged *C*_*E*_. Bioelectric fields, morphogenetic patterns, and related coordination substrates (Levin, 2019) operate at the ensemble level above *C*^∗^, not at microstate trajectory resolution; the framework locates their role, it does not deny it.

### 2.9 Computational verification

We tested the same threshold structure in three complementary computational model classes: AlphaFold-2 cross-organism proteomes, the JCVI-syn3A 4D whole-cell simulation, and four canonical stochastic gene-network architectures. AlphaFold and syn3A are large established models built without reference to the present framework, while the stochastic-network simulations test whether the same coarse-to-fine support growth recurs across unrelated subcellular dynamical modules. Together, the three analyses span molecular, cellular, and module-level scales.

#### 2.9.1 AlphaFold cross-organism folding gap

We aggregated structural information across 46 AlphaFold-2 proteomes spanning archaea, bacteria, fungi, amoebozoa, invertebrates, plants, and vertebrates (555,846 proteins, 241,282,135 residues) (Jumper *et al*., 2021; Varadi *et al*., 2022). For each protein we computed a 1 Å structural-bit estimate *B*_1Å_= 3*N*_atom_ log_2_(*L/*1Å). Coding-input bits were *B*_code_ = 6*L* per protein (the mechanistically relevant input to folding); haploid whole-genome bits *B*_hap_ = 2 × bp were the strongest organism-specific storage upper bound. The scientifically meaningful comparison charges the shared compiler cost *H*(*M*) exactly once across the panel: *R*_agg_(*D*) = ∑_*i*_ *B*_1Å,*i*_*/*(*H*(*M*) + ∑_*i*_ *D*_*i*_), with *H*(*M*) ≈ 3.0 × 10^9^ bits at fp32 (Jumper *et al*. 2021; Methods).

Aggregate structural output (38.5 Gb) exceeds aggregate coding input (1.4 Gb) by ∼27×; after paying once for the shared model, the ratio is 9–21× (Table 2, Fig. 2). The proteome/exome ratio is approximately constant at ∼21–30× across the entire 46-organism panel. The proteome/genome ratio, by contrast, varies by nearly two orders of magnitude, descending from ∼20× in bacteria to ∼0.4× in vertebrates. If extra-exomic DNA carried the folding bits, the proteome/genome ratio should be the more stable of the two, since genome size includes both coding and non-coding storage; the empirical pattern is the opposite, which is what one expects if a single compiler operates on per-protein coding input and not on lineage-specific non-coding storage.

**Table 2:** Aggregate 46-organism information balance for the AlphaFold folding map. Ratios above 1 indicate structural output exceeds the input budget.

| Denominator | Aggregate bits (Gb) | $B_{1\text{\AA}}/D$ | $B_{1\text{\AA}}/(D + H(M))$ |
| --- | --- | --- | --- |
| Coding input $B_{\text{code}}$ | 1.448 | 26.58 | 8.65 (fp32), 21.11 (4-bit) |
| Haploid genome $B_{\text{hap}}$ | 30.9 | 1.244 | 1.134 (fp32), 1.229 (4-bit) |

**Figure 2:**
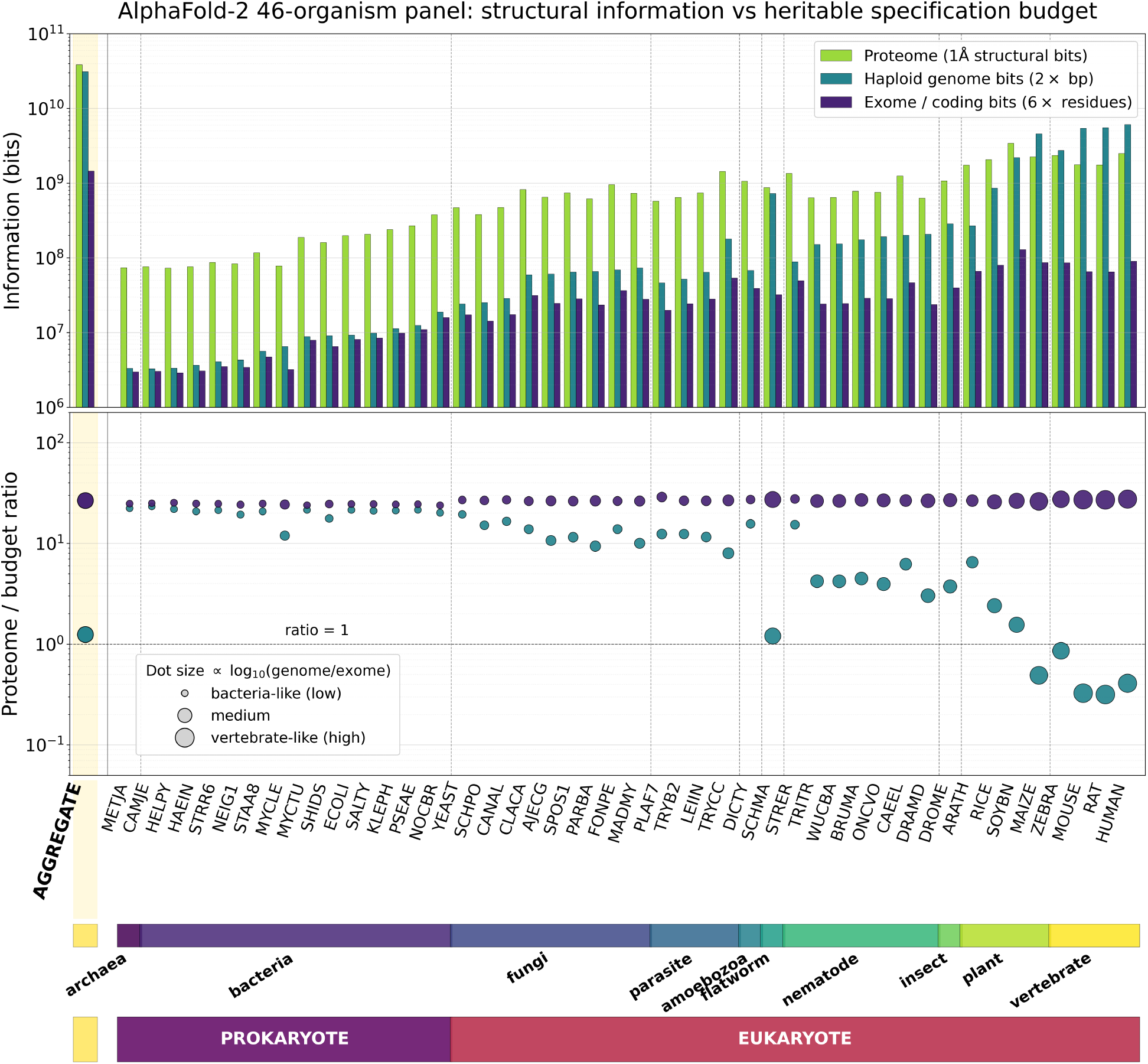
Structural output exceeds heritable specification budget across 46 organisms. **Top:** per-organism proteome structural bits (yellow-green), haploid genome (teal), and exome (purple). **Bottom:** corresponding ratios. The proteome/exome ratio is approximately constant (∼21–30×) across all 46 organisms despite genomes spanning four orders of magnitude. The proteome/genome ratio varies by nearly two orders of magnitude, descending from ∼20× in bacteria to ∼0.4× in vertebrates (the C-value paradox made quantitative).

The same conclusion holds for the MSA-free predictor ESMFold (Lin *et al*., 2023), whose ∼ 1.5×10^10^ parameters amortize to only ∼780 bits/protein over ∼ 6.17 × 10^8^ predictions. Any residual extra-exomic effect on folding must travel through the current cellular state, where, by Lemma 2, it counts against the same causal-locality budget as current function.

Across the 241,282,135 residues in the 46-organism panel, the idealized backbone-only *ϕ/ψ* description equals the aggregate coding input of 1.448 Gb at a resolution of 45^°^ per torsion and exceeds it at every finer resolution (Supplementary Table S2).

#### 2.9.2 JCVI-syn3A 4D whole-cell simulation

We computed information costs across a coarse-graining ladder on the publicly deposited 4D simulation of *Mycoplasma* JCVI-syn3A (Luthey-Schulten *et al*., 2026), the most detailed whole-cellcycle simulation published to date. We sampled 13 frames at 600-second intervals from each of four complete cell cycles (MinCell 1–4) over *t* = 0 to 7200 s, for 52 snapshots in total. The syn3A genome is 543,379 bp, giving a raw budget *C*_*G*_ ≈ 1.087 × 10^6^ bits at two bits per nucleotide. For each snapshot we computed the conservative spatial zero-error addressability cost 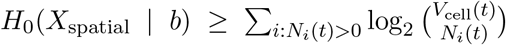, treating within-species particles as indistinguishable over *V*_cell_(*t*) occupied lattice sites; this bound deliberately undercounts the true addressability cost (Methods).

Three results are reported in Table 3 and Fig. 3. First, the conservative unlabeled-position bound on *H*_0_(*X*_spatial_ | *b*) already exceeds the genomic budget at full biomass (139% of *C*_*G*_ at *t* = 7200 s). Second, the spatial zero-error addressability cost grows monotonically with biomass, from ∼60% of *C*_*G*_ at *t* = 0 to 139% at *t* = 7200 s, crossing *C*_*G*_ at median *t* ≈ 4300 s across the four trajectories (per-trajectory crossings 4200–4840 s; Fig. 3b). Third, copy-number and per-compartment descriptors consume only 0.31% and 1.1% of *C*_*G*_ respectively, while a trajectory-level descriptor under a 10-second decorrelation choice reaches ∼ 665×*C*_*G*_. Functional descriptors are two orders of magnitude below the genomic budget; trajectory descriptors are three orders of magnitude above it. The threshold *C*^∗^ sits between the compartment-level and per-snapshot spatial descriptors, and is crossed dynamically during the cell cycle.

**Table 3:** Information ladder for the JCVI-syn3A 4D whole-cell simulation. Spatial and trajectory rows (*C*_spatial_, *C*_trajectory_) report zero-error addressability quantities *H*_0_(*X* | *b*). The compartment-level row reports a marginal NSB Shannon estimate. The trajectory envelope is heuristic and depends on the assumed decorrelation time. The raw haploid genomic budget *C*_*G*_ is 1,086,758 bits.

| Descriptor | Information quantity (bits) | Ratio to $C_G$ |
| --- | --- | --- |
| $\mathcal{C}_{\text{prot}}, \mathcal{C}_{\text{joint}}$ (copy number) | 3,353/3,357 | 0.31% |
| $\mathcal{C}_{\text{compartment}}$ (per-region per-species, NSB upper) | 12,116 | 1.11% |
| $\mathcal{C}_{\text{spatial}}$ ($t = 0$, lower bound) | 656,321 | 60% |
| $\mathcal{C}_{\text{spatial}}$ (median across snapshots, lower bound) | 1,003,926 | 92% |
| $\mathcal{C}_{\text{spatial}}$ ($t = 7200$ s, lower bound) | **1,506,981** | **139%** |
| $\mathcal{C}_{\text{trajectory}}$ ($T_{\text{eff}} = 720$, 10 s decorrelation, 2 h) | $7.23 \times 10^8$ | 66,500% |
| $C_G$ (syn3A raw haploid genome) | 1,086,758 | 100% |

**Figure 3:**
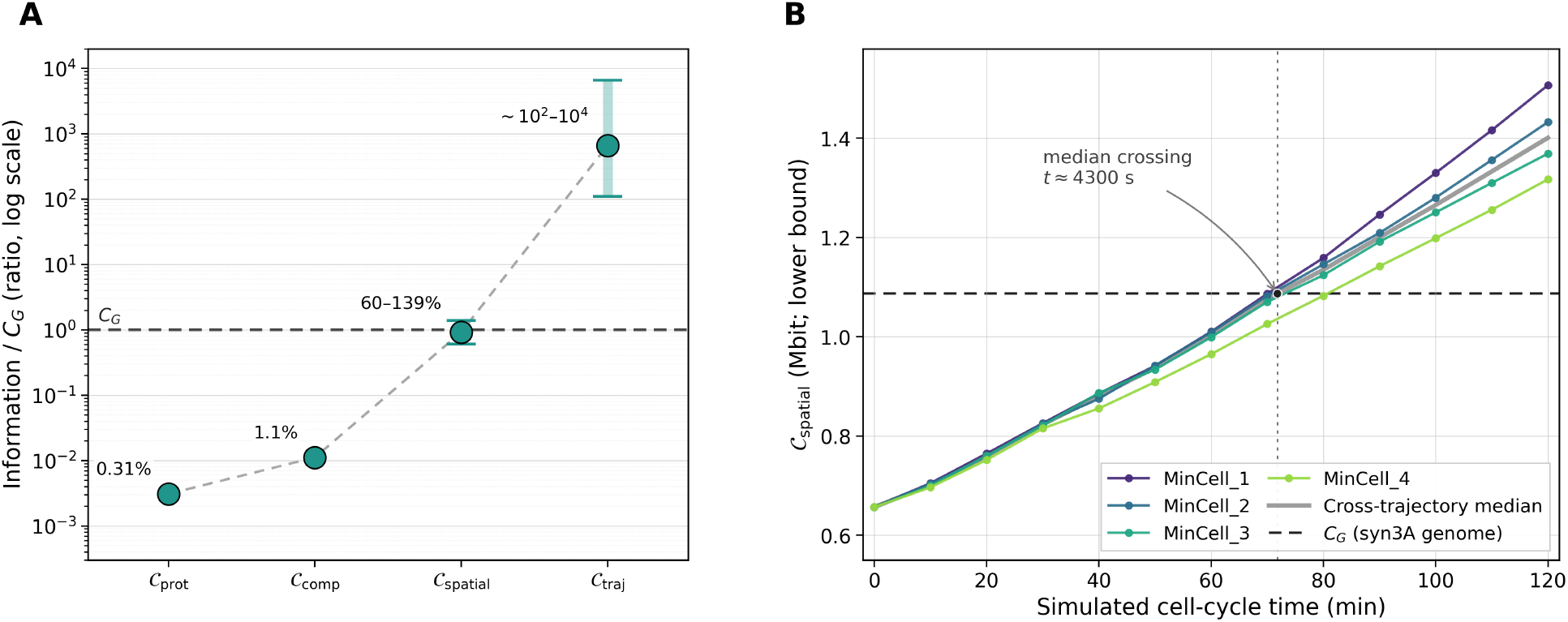
Threshold structure in the JCVI-syn3A 4D whole-cell simulation. **(a)** Information ladder across coarse-grainings, expressed as ratio to the syn3A genomic budget *C*_*G*_ ≈ 1.087 Mbit. The total range across the ladder spans approximately six orders of magnitude. **(b)** Spatial zero-error addressability cost over simulated time for four cell cycles (MinCell 1–4). All four trajectories cross *C*_*G*_ between *t* ≈ 4200 and 4840 s (median ≈ 4300 s), approximately mid-cell-cycle.

#### 2.9.3 Canonical stochastic network simulations

We tested whether the coarse-to-fine support growth observed at the whole-cell level also appears within individual sub-cellular modules. We simulated four canonical stochastic architectures spanning distinct dynamical regimes — a two-state bursting gene-expression model (Peccoud & Ycart, 1995), the Elowitz–Leibler repressilator (Elowitz & Leibler, 2000), the Gardner–Cantor–Collins toggle switch (Gardner *et al*., 2000), and a simplified MAPK cascade with feedback after Munsky *et al*. (2012) — using the Gillespie algorithm across five seeds per model. Each trajectory was projected onto a coarse-graining ladder from protein-count projections (*C*_prot_) to the full biochemical count state (*C*_joint_) to post-hoc unlabeled occupancy refinements (*C*_occ2d_(*g*), a conservative combinatorial localization descriptor, not a reaction–diffusion simulation).

Across all four architectures, biochemical coarse-grainings remain compact (8–16 bits per network) and localization-like refinements rapidly expand the observed support (Fig. 4b). In unsaturated rows (*K*_obs_*/N <* 0.8) we report NSB Shannon estimates; in saturated rows, nearly every snapshot maps to a distinct symbol, and the empirical result is the support-growth signal itself. The same hierarchy — compact biochemical descriptions, rapidly saturating fine localization — recurs across all four unrelated architectures.

**Figure 4:**
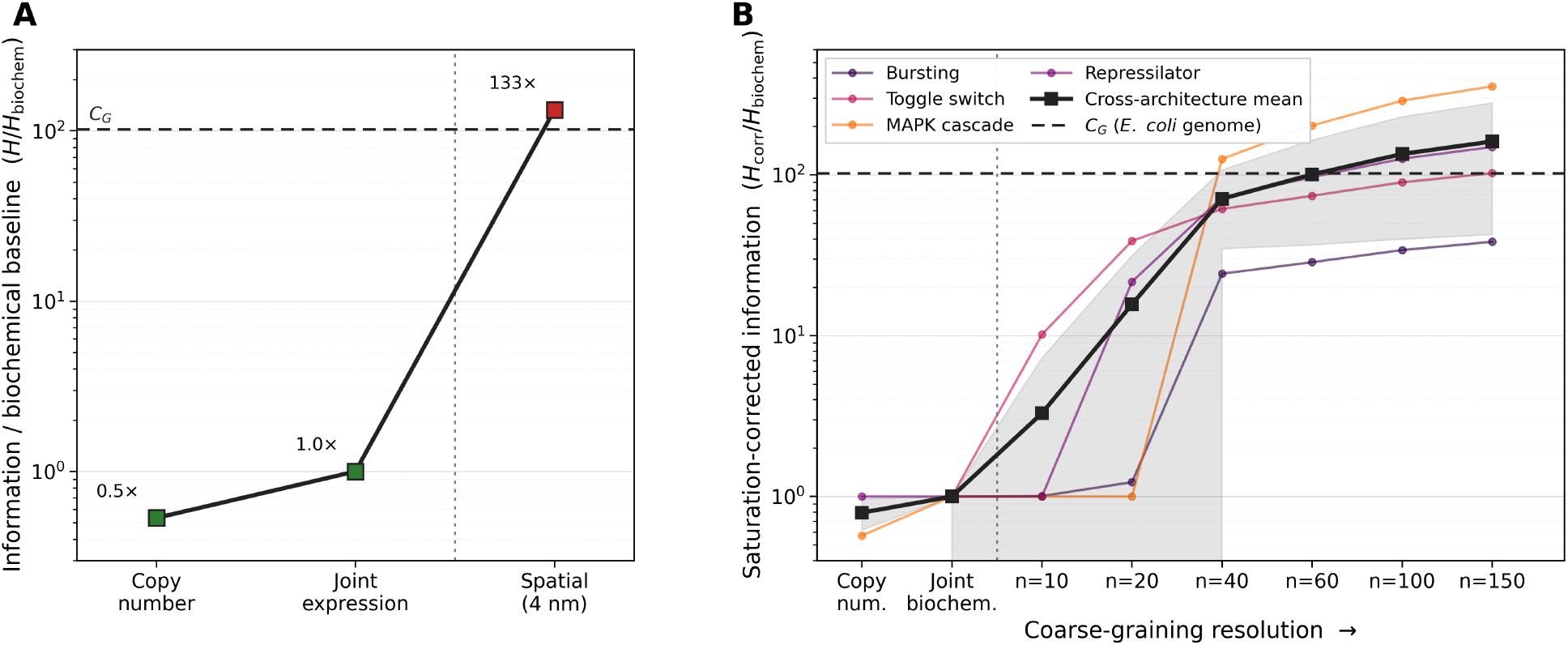
Information versus coarse-graining resolution in whole-cell and module-level examples. **(a)** Whole-cell threshold crossing for *E. coli*. Functional copy-number and jointexpression descriptors remain below the raw genomic budget, whereas a conservative 4 nm spatial addressability descriptor crosses it. The spatial row is an *H*_0_ zero-error addressability quantity, not a Shannon entropy estimate. **(b)** Module-level corroboration across four canonical stochastic architectures. Thin lines show individual models; the bold black line shows the cross-architecture mean. Unsaturated biochemical rows report empirical NSB Shannon estimates. Saturated occupancy rows are support-envelope quantities and indicate that the localization descriptor support has exceeded finite sampling.

## 3 Discussion

In this study, we establish information-theoretic limits on the programmatic specification of biological systems. We demonstrate that an organism lacks the information capacity to specify its own fully functioning microscopic organization. By formalizing a coarse-graining information threshold, we show that organism-controlled information — the genome and continuous environmental signals — is sufficient to specify macroscopic functional ensembles, but falls short of programming exact molecular trajectories by orders of magnitude. We located this limit empirically across *E. coli, Drosophila*, and *C. elegans*, and verified its predicted structure in state-of-the-art computational models (AlphaFold, the JCVI-syn3A whole-cell simulation, and canonical stochastic networks). Finite genomic budgets comfortably accommodate cell fates, connectomes, and compartments, but are vastly exceeded by exact spatial coordinates. We do not claim that organism-controlled information cannot encode compact, functionally relevant summaries of microstates that it cannot fully specify. A recent computational model demonstrates that topology- and function-associated protein organization can be retained in a compressed latent code (Guo *et al*., 2026); such summaries are not the physical macromolecular organization itself, whose fine-grained programmatic specification our bounds exclude.

Consequently, the genome must be understood as a blueprint of an ensemble-level generator, not a microscopic trajectory program. It specifies molecules, regulatory architecture, boundary conditions, and interaction rules whose physical evaluation produces a distribution of admissible outcomes. Non-DNA coordination substrates — such as bioelectric fields, mechanical patterning, and morphogenetic waves (Levin, 2019) — operate at this same ensemble level. They synchronize, bias, and constrain developmental generators rather than specifying microstate trajectories; attempting the latter would violate the same addressability, causal-locality, and channel-capacity limits as DNA.

Crucially, this framework recasts biological specification as a form of lossy compression. The genome is a compact organism-specific message, and the physical substrate acts as an active universal compiler, decoding this message into an ensemble of admissible realizations. Because this compiler sits outside the specification budget, the genome cannot fully control the microstate residual. Instead, it biases the compiler through sequences and regulatory rules, while physical dynamics actively resolve the remaining microscopic degrees of freedom. Biological function must therefore be organized around invariants that survive this compiler-induced residual, rather than relying on exact microstate identity.

This distinction also clarifies what belongs to the environmental specification capacity *C*_*E*_. The total physical environment cannot be identified with *C*_*E*_: a suitable substrate can be essential without specifying organismal identity. Many different seeds can be planted in the same soil, and the soil alone does not determine which plant will develop. The seed carries the organism-specific generator specification; the soil, temperature, water, and chemistry provide the permissive substrate in which that specification is compiled. Only signals that carry decodable information about the particular organismal outcome belong to *C*_*E*_; raw environmental complexity is not organism-specific specification.

This structural realization yields four derivative consequences for the study of biological systems. First, the phenotype space accessible to evolutionary search is strictly the ensemble of generator outputs. A genome cannot contain a complete microstate-resolution genotype-to-phenotype simulator; the theorem mathematically rules out such strong internal forward models. Rather, the genome encodes coarse forward biases — developmental constraints, modularity, and canalization — that make certain phenotypic regions more accessible. This explains how evolution discovers complex physical optimizations, such as the local branching geometry of vasculature or plant structures (Meng *et al*., 2026). Evolution does not search infinite microscopic coordinate spaces; the genome supplies sparse parameters, and shared surface-minimization physics computes the optimized geometry. Selection acts retrospectively on realized macroscopic statistics, shifting genotype frequencies without ever encoding the full trajectory.

Second, the theorem clarifies the mathematical relationship between heritability and genetic determinism. Heritability measures how strongly genetic differences account for variation in a chosen coarse phenotype (Falconer & Mackay, 1996; Visscher *et al*., 2008). However, a shared and stable physical compiler may be causally indispensable while contributing negligible between-individual variance, rendering it largely invisible to heritability estimates. Thus, a genotype may predict a coarse phenotype with high heritability, even though the exact microscopic realization retains residual entropy and remains genetically unspecified. Heritability represents variation around a shared compilation process, not evidence for a complete genetic program.

Third, our results refine thermodynamic accounts of life. Schrödinger framed biological order as requiring free-energy throughput and inherited information (Schrödinger, 1944). The threshold theorem defines this inherited information precisely — a coarse generator specification — and partitions the remaining order. Free energy maintains the nonequilibrium conditions for compilation, while the physical substrate supplies the shared regularities (e.g., folding and diffusion) that execute it. Biological order is thus the product of a three-way division of labor: organism-specific coarse specification, nonequilibrium maintenance, and shared physical compilation. This division motivates a quantitative bridge to stochastic-thermodynamic accounts of information flow (Seifert, 2012; Horowitz & Esposito, 2014), suggesting testable signatures in dissipation and error correction.

Fourth, our work sets absolute limits on organism-internal specification for computational modeling and engineering. Any predictive model operating below the coarse-graining threshold must approximate a compiler-induced input–output map rather than recover an internal blueprint. This explains the cross-organism generalization of generative models such as AlphaFold, ESMFold, and Evo (Jumper *et al*., 2021; Lin *et al*., 2023; Nguyen *et al*., 2024): they approximate shared physical, biochemical, or evolutionary structures, not lineage-specific microscopic programs. Mechanistic interpretation can recover compiler features, but not an internal master program tape, because no such tape exists. Similarly, engineered interventions such as CRISPR and optogenetics are bounded by these capacities; they bias pathways and specify ensemble behaviors, but cannot script exact downstream molecular trajectories.

While minimal systems with small state spaces — such as viral capsids and isolated catalytic complexes — may approach microscopic resolution, complex biology necessarily enters the ensemble regime at sufficiently fine resolution. Furthermore, while this framework intersects several adjacent literatures, it addresses a distinct question. Classical zero-error capacity concerns channel-specific addressability (Shannon, 1956); Mori–Zwanzig projection methods derive reduced macrodynamics from microscopic dynamics (Zwanzig, 1961; Mori, 1965); and stochastic thermodynamics under coarse-graining quantifies hidden entropy production (Esposito, 2012; Kawaguchi & Nakayama, 2013). We instead quantify what an organism-controlled specification can program. The information measure itself is transmission information (Shannon, 1948; Cover & Thomas, 2006); biological meaning enters through function, via the coarse-grainings at which biological functions are empirically defined.

In summary, the threshold theorem establishes the genome as a generator specifier by ruling out organism-programmed microstate determinism while leaving physical determinism untouched. The result reframes specification, modeling, and engineering of biological systems around a single inequality, and warrants substantial follow-up work across information theory, thermodynamics, evolutionary theory, and synthetic biology. The genome specifies the generator. The universal compiler, operating on the physics substrate, computes the samples. Selection acts on the statistics of the resulting ensemble.

## 4 Methods

### 4.1 Entropy estimators

All entropy estimates are reported in bits. For an integer count vector *n* = (*n*_1_, …, *n*_*K*_) with total sample size *N* = ∑_*i*_ *n*_*i*_ over an observed alphabet of s ize *K*_obs_ = |{*i* : *n*_*i*_ *>* 0}|, we used three estimators in parallel: the plug-in estimator 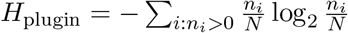, the Miller–Madow bias-corrected estimator 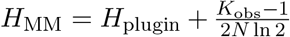, and the Nemenman–Shafee–Bialek (NSB) estimator, which we used as the primary estimator throughout because it is well-behaved in the undersampled regime characteristic of stochastic-network and whole-cell-simulation data (Nemenman *et al*., 2002).

The NSB posterior mean was evaluated by integrating over a symmetric Dirichlet concentration parameter *β* on a 200-point logarithmic grid spanning log *β* ∈ [−8, 8]. For a hypothesized total alphabet size *K*, the per-*β* log evidence is log *p*(*n* | *β*) = log Γ(*Kβ*) −log Γ(*Kβ* + *N*) + ∑ log Γ(*β* + *n*_*i*_)−log Γ(*β*), and the posterior expected entropy in nats is 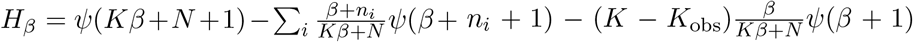 where *ψ* denotes the digamma function. The NSB prior was implemented through the standard uniform-on-entropy reparameterization Jacobian *dξ/dβ* = *Kψ*^*′*^(*Kβ* + 1) − *ψ*^*′*^(*β* + 1), with nonpositive numerical values clipped to zero. When a physical or combinatorial upper bound *K*_upper_ on the alphabet size was available, we enforced *K*_NSB_ = max{*K*_upper_, *K*_obs_, 2} to avoid the ill-defined regime *K < K*_obs_.

We treated rows with *K*_obs_*/N* ≥ 0.8 as saturated. In this regime the plug-in and Miller–Madow estimators are sample-size limited, and the NSB estimate is best interpreted as a *K*_upper_-conditioned upper envelope rather than a sample-driven point estimate. Saturation rows are flagged as such throughout the figures and tables.

### 4.2 Stochastic network simulations

We simulated four canonical stochastic biochemical networks using the Gillespie direct method(Gillespie, 1977). The state vector 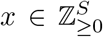, propensity vector *a*(*x*), and stoichiometry *ν*_*r*_ were specified per model; at each step we drew Δ*t* ∼ Exp(∑ _*r*_ *a*_*r*_(*x*)) and a reaction index proportional to *a*_*r*_(*x*), and updated *x* ← *x* + *ν*_*r*_. To remove the dwell-time bias inherent in event-resolved trajectories, we discarded the initial 20% of reactions as burn-in and resampled each trajectory as a right-continuous piecewise-constant path at uniform wall-clock spacing.

#### Model specifications

The four networks were the two-state bursting gene-expression model (Peccoud & Ycart, 1995), the Elowitz–Leibler repressilator (Elowitz & Leibler, 2000), the Gardner– Cantor–Collins toggle switch (Gardner *et al*., 2000), and a simplified three-tier MAPK signaling cascade with feedback after Munsky *et al*. (2012). Parameter values were taken from measured *E. coli* single-molecule distributions (Taniguchi *et al*., 2010) for the bursting model, and from the original publications for the others. Bursting parameters: *k*_on_ = 0.5, *k*_off_ = 1.0, *k*_tx_ = 20.0, *γ*_*m*_ = 1.0, *k*_tl_ = 5.0, *γ*_*p*_ = 0.1. Toggle switch: *α*_1_ = *α*_2_ = 40.0, *β* = 3.0, *γ* = 4.0, *K* = 30.0, degradation rate 1.0. Repressilator: *α* = 50.0, *α*_0_ = 0.5, *K* = 40.0, Hill coefficient *n* = 2.0, *γ*_*m*_ = 1.0, *k*_tl_ = 5.0, *γ*_*p*_ = 0.2. MAPK cascade: 10 Michaelis–Menten reactions across the three phosphorylation tiers, with conservation laws total MKKK = 100, total MKK = 300, total MAPK = 300. Each model was run with five independent random seeds. Production runs used either 10^6^ or 3 × 10^6^ maximum reactions per trajectory and uniform resampling intervals of 1.0 s or 0.2 s; the canonical-network figure uses the 3 × 10^6^, 0.2 s configuration.

#### Coarse-graining ladder

Each resampled trajectory was projected onto three coarse-grainings of increasing fineness:

- *C*_prot_: protein-count projection, retaining only protein-like species counts.
- *C*_joint_: the full biochemical count vector across all tracked species.
- *C*_occ2d_(*g*): a post-hoc unlabeled hard-core occupancy descriptor on a *g* × *g* lattice, appended to the biochemical state vector. For a snapshot with total molecule count *n*_tot_ ≤ *g*^2^, the descriptor draws *n*_tot_ distinct lattice cells uniformly at random without replacement and appends their sorted indices. For *n*_tot_ *> g*^2^, the descriptor appends an overflow sentinel and is degenerate with *C*_joint_ plus a constant marker. This descriptor is intentionally a conservative combinatorial localization refinement, not a mechanistic spatial reaction–diffusion simulation, and the framework’s claim is that even this conservative refinement triggers the entropy explosion. We swept *g* ∈ {10, 20, 40, 60, 100, 150}.

Overflow rows were reported separately and not interpreted as spatial threshold crossings. Saturated non-overflow rows were interpreted as direct evidence that the effective support exceeds the finite sample budget — that is, the state space at that resolution is too large to be exhaustively sampled. A pilot bounded-support moment descriptor was tested but retired: when random positions are drawn independently per snapshot it injects Monte Carlo noise into the estimate, and when the random generator is deterministically seeded by the biochemical state it collapses to a relabeling of *C*_joint_. This null result is reported in the Supplementary Material and is not used for any main-text claim.

### 4.3 AlphaFold cross-organism structural accounting

We analyzed 46 AlphaFold-2 proteomes downloaded from the EBI AlphaFold public FTP (Varadi *et al*., 2022). The panel comprises 16 model organisms spanning archaea, bacteria, fungi, amoebozoa, invertebrates, plants, and vertebrates, plus 30 Global Health proteomes covering bacterial pathogens, parasites, pathogenic fungi, and nematodes; the full organism list and source URLs are provided in the Supplementary Material. For each proteome, we parsed every compressed mmCIF file in the archive (no protein-level subsampling), reading heavy atoms (excluding hydrogens) from`_atom_site` fields, recording Cartesian coordinates, accumulating pLDDT confidence values, and tracking unique residue indices.

#### Structural information at 1Å geometric resolution

For each organism, the structural information content at 1Å geometric resolution was estimated as *B*_1Å_= 3*N*_atom_ log_2_(*L/*1Å), where *N*_atom_ is the total heavy-atom count in the proteome and *L* is the median, over proteins, of the maximum coordinate-axis span plus a 10Å padding term to accommodate atomic excursions beyond the bounding box. Sensitivity to the resolution choice was tested at 0.5Å and 2.0Å, and is reported in the Supplementary Material. The factor of three reflects the three Cartesian coordinates per atom; the logarithm bounds the bits required to specify each coordinate at the chosen resolution within the bounding box. The 1 Å scale is the relevant functional-addressability resolution for protein structure: many biological functions — binding pockets, catalytic residues, steric compatibility, hydrogen bonds, salt bridges, and docking interfaces — depend on atomic-scale contact geometry that is not specified by global fold class or low-dimensional shape descriptors alone.

#### Genomic and coding denominators

Genome size in base pairs was taken from the corresponding NCBI Assembly entries. Haploid bits were computed as *B*_hap_ = 2×(haploid genome size in bp), reflecting two bits per nucleotide for the four-letter DNA alphabet. As established in the framework section, this haploid bit count is also a tight upper bound on the diploid information content (within ∼ 1% via the heterozygosity inequality), so we did not separately compute a diploid denominator. The coding denominator was *B*_code_ = 6 ∑_*i*_ *L*_*i*_, using three nucleotides per residue at two bits per nucleotide.

#### Aggregate compiler-amortized comparison

The scientifically meaningful comparison is the aggregate ratio 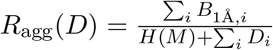, which charges the shared compiler cost *H*(*M*) exactly once across the panel. For the AlphaFold-2 model size we used *H*(*M*) ≈ 3.0 × 10^9^ bits at fp32 precision and ∼ 3.75 × 10^8^ bits at 4-bit precision, with intermediate fp16/bf16 (∼ 1.49 × 10^9^ bits) and 8-bit (∼ 7.5 × 10^8^ bits) values reported in the Supplementary Material as sensitivity points. These model-bit terms are used solely to test the literal lookup-table interpretation of the opaque learned predictor; auxiliary external inference information (MSAs, structural templates, database-derived context) is treated as predictor-side input and is not included in the per-organism specification budget *D*_*i*_, since it is neither present in the folding cell nor part of the organism’s heritable store.

#### ESMFold sequence-only comparison

For the MSA-free corroboration, we used the published ESMFold parameter count of ∼ 1.5 × 10^10^ parameters (Lin *et al*., 2023), corresponding to *H*(*M*_ESM_) ≈ 4.8×10^11^ bits at fp32, and the published ESM Metagenomic Atlas scale of ∼ 6.17×10^8^ predicted structures with ∼ 2.25 × 10^8^ at high confidence. The structural-output scale per protein was estimated from the same 46-organism panel as 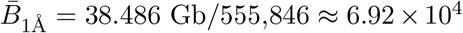 bits per protein. The amortized per-protein model cost ∼ 780 bits over the full Atlas (or ∼ 2100 bits over the high-confidence subset) is the relevant comparison: even with no MSA at inference, the model is far smaller than would be required for a per-protein structural lookup table.

### 4.4 JCVI-syn3A 4D whole-cell simulation analysis

We analyzed the publicly deposited JCVI-syn3A 4D whole-cell simulation data of Luthey-Schulten *et al*. (2026), available as `Minimal_Cell_4DWCM.zip` on Zenodo. The archive contains four complete Lattice Microbes simulated cell cycles (`MinCell_1.lm` through `MinCell_4.lm`) and 50 coarse count/flux trajectories. Our analysis used the four full-state HDF5 trajectories.

Each `.lm` file contains lattice frames of shape 128 × 64 × 64 × 16 with 10 nm lattice spacing, 5489 tracked molecular species, seven site types (extracellular, cytoplasm, outer cytoplasm, ribosomes, ribosome centers, DNA, and membrane), and 16 slots per lattice site. Frame indices correspond to wall-clock seconds. We sampled 13 frames per trajectory at 600-second intervals over 0–7200 s, yielding 52 snapshots in total.

#### Genomic budget

The syn3A genome is 543,379 bp; we used *C*_*G*_ = 543,379 × 2 = 1,086,758 bits as the maximally generous raw genomic budget, computed at two bits per nucleotide for the four-letter DNA alphabet. The factor of two is the bp-to-bits conversion, not a diploidy adjustment; the simulation models a bacterial cell whose two genome copies (during cell cycle stages with replicated chromosomes) are clonally identical and therefore do not carry independent information.

#### Spatial zero-error addressability cost

For each snapshot, the conservative spatial zero-error addressability cost *H*_0_(*X*_spatial_ | *z*) was computed by pooling counts across intracellular regions and treating particles within each species as indistinguishable over *V*_cell_(*t*) occupied cellular lattice sites: 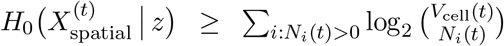, where *N*_*i*_(*t*) is the count of species *i* at time *t*. The right-hand side bounds the number of distinguishable spatial configurations consistent with the snapshot. This descriptor deliberately undercounts the simulation state in three ways: it ignores within-species particle labels, it ignores the 16-slot ordering within each lattice site, and it does not include conformational, modification, or binding-state information. The actual zero-error addressability cost is therefore strictly larger than this bound, so any reported threshold crossing under Theorem 1 part 2(a) is conservative in the direction that matters.

#### Compartment-level Shannon estimate

The compartment-level descriptor *C*_compartment_ was computed by binning species occupancy across the seven site types, building a per-(region, species) count matrix across snapshots, and summing marginal NSB entropies under the marginal-independence assumption. This is an upper bound on Shannon entropy rather than on *H*_0_: the marginalindependence assumption ignores cross-species correlations that would only reduce the joint Shannon entropy.

#### Trajectory envelope

A trajectory-level addressability envelope was estimated as 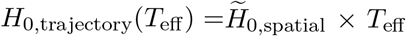, ^where 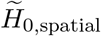^ is the median spatial-snapshot addressability cost and *T*_eff_ is the effective number of independent timepoints over the 2-hour window. We reported *T*_eff_ ∈ {120, 720, 7200}, corresponding to one independent snapshot every 60, 10, or 1 second respectively. The choice of *T*_eff_ is heuristic and depends on the assumed decorrelation time; the inequality direction (trajectory addressability cost ≫ *C*_*G*_) is robust to this choice across more than two orders of magnitude.

### 4.5 Bicoid information channel and continuous-environmental rate calculations

The Bicoid–Hunchback channel-capacity values reported in the main text are taken from published quantitative measurements: ∼ 1.5–1.7 bits per nucleus per fresh Bicoid readout, nuclear recovery time *τ*_*c*_ = 68.9±17.6 s, and ∼ 4.3±0.1 bits for the four-gap-gene positional code (Gregor *et al*., 2007; Tkačik *et al*., 2008; Petkova *et al*., 2019). The integrated continuous-environmental information bound used the standard sampling expression *I*(*X*; *U*_[0,*T*]_) ≤ ⌈*T/τ*_*c*_⌉*I*_∗_, applied to the one-hour developmental window. The resulting per-nucleus environmental budget of ∼ 80–230 bits depends on whether one adopts the Bicoid–Hunchback channel alone or the four-gap-gene positional code as *I*_∗_.

The supplementary modality-specific calculations for chemical, mechanical, electrical, electromagnetic, thermal, and quantum signaling channels (Supplementary Note 2) use explicit source, receiver, rate, fidelity, addressability, synchronization, and actuation assumptions. Chemical signaling was treated using Berg–Purcell counting statistics and a Gaussian-equivalent information-rate proxy under assumptions deliberately generous to the rescue hypothesis; mechanical, electrical, electromagnetic, and thermal channels were treated as parameter-conditioned physical envelopes rather than amplitude-independent universal capacities. Quantum signaling was treated using Holevo information accounting and explicit source, coherence, receiver, and actuation requirements; no universal cytoplasmic decoherence-rate capacity bound was assumed. The purpose is not to deny that environments shape biological ensembles — they clearly do — but to test whether any known naturally realized source–receiver architecture can act as a deterministic, addressable microstate instruction tape. Engineered external controllers are not ruled out; their added sensing, memory, communication, and actuation capacity belongs to an enlarged *C*_*E*_.

### 4.6 Data and code availability

All source code, configuration files, parameter specifications, and analysis scripts used to reproduce the computational-model results and the figures and tables derived from them are available at https://github.com/yuj1r0/itol. Large public input datasets are not redistributed but are reproducible from their original sources: AlphaFold-2 proteomes from the EBI AlphaFold public FTP (Varadi *et al*., 2022), the JCVI-syn3A 4D whole-cell simulation data from Zenodo record 15579159 (Luthey-Schulten *et al*., 2026), and the NCBI Assembly genome size annotations from the corresponding accessions. Stochastic network simulation outputs are reproducible from the configuration files and random seeds in the repository. An end-to-end reproduction recipe is included.

