## Supplementary Material for "Information-theoretic Limits on Programmatic Specification of Biological Systems"

July 27, 2026

This supplement contains two notes. **Supplementary Note 1** (Extended Mathematical Framework) gives the formal physical framework, the formal definitions of coarse-graining and programmed microstate determinism, the core lemmas, the zero-error addressability proposition, and the main coarse-graining threshold theorem in full, with complete statements and proofs, followed by appendices providing precise information-theoretic notation, the underlying probability space, the connection between the runtime-randomness lemma and continuous-time Markov dynamics, the relationship between Hartley and Shannon capacity bounds, the contraction assumption underlying the mixing lemma, and a precise restatement of the causal-locality lemma. **Supplementary Note 2** (Environmental Rescue Bounds) develops the modality-specific bounds summarized in the main text, with explicit rate, addressability, and feedback-vs-override analyses for chemical, mechanical, electrical, electromagnetic, thermal, and quantum signaling, followed by direct replies to common objections concerning environmental richness, omitted channels, alternative substrates, and inherited oocyte or cytoplasmic state.

### Supplementary Note 1: Extended Mathematical Framework

#### 1 Formal physical framework

To rigorously establish information-theoretic limits on biological specification, we formalize the organism as a physical information-processing system. We model the biological system not merely as a finite developmental endpoint, but as a continuous-time physical computation running over an arbitrary temporal window $[t_0, t_n]$. Analogous to an operating system computing a trajectory of states based on a static executable and a continuous stream of hardware interrupts, the biological system computes a physical microstate trajectory based on a static genomic specification and a continuous stream of environmental inputs.

We define the system and its available information stores through the following formal objects (Fig. 1):

1. **Microstate space ($\Sigma$) and physical trajectory:** Let $\Sigma$ denote the physical state space at the resolution of full molecular microstate. At any instant $t$, the state $S_t \in \Sigma$ captures the exact spatial coordinates, conformational states, modification statuses, and binding partners

of every molecule within the system boundaries. Because the system comprises a bounded number of molecules in a finite volume and spatial and conformational coordinates are resolved at some chosen physical (e.g., Å-scale) resolution, $\Sigma$ is finite; equivalently, $\Sigma$ is the discretization, at that resolution, of the compact configuration manifold of the finite molecular system, and therefore has finite cardinality.

We fix a time-discretization $t_0 = \tau_0 < \tau_1 < \cdots < \tau_K = t_n$ at resolution $\delta = \tau_{i+1} - \tau_i$. The realized physical outcome over the temporal window is the discretized trajectory $\mathbf{S} = (S_{\tau_0}, S_{\tau_1}, \ldots, S_{\tau_K}) \in \Sigma^{K+1}$. All entropies in this paper refer to finite random vectors of this form. The choice of $\delta$ is a modeling parameter; all impossibility results hold for every $\delta > 0$, and in particular for the finest resolutions used in the computational verifications. The underlying probability space and the relationship between this discrete framework and the continuous-time Markov dynamics of the physical system are described in Appendix B.

2. **Genomic channel ($g$):** The static, onboard, heritable information store. A genome of $n$ base pairs is a string over a four-letter alphabet; we define its specification capacity as the Hartley entropy $C_G = \log_2 4^n = 2n$, i.e., the logarithm of the number of possible genome strings of that length. This is an upper bound on the Shannon entropy $H(g)$ for any distribution over genomes, and it is the quantity computed in all empirical comparisons below. In empirical comparisons we use the full raw haploid sequence as a deliberately generous upper bound on organism-specific genomic capacity. Functional capacity can only be smaller, so any threshold crossing against this budget is conservative and does not depend on classifying sequence as coding, regulatory, or inert. For the Hartley prong of the main theorem, $C_G$ bounds the number of distinguishable messages the genome can carry; for the Shannon prong, it upper-bounds $H(g)$ (see Appendix D).

3. **Continuous environmental channel ($U_{t_0:t_n}$):** The integrated stream of exogenous physical inputs (chemical, mechanical, electromagnetic, thermal) crossing the system boundary during the interval. Under the time-discretization above, $U_{t_0:t_n}$ denotes the finite sequence $(U_{\tau_0}, U_{\tau_1}, \ldots, U_{\tau_{K-1}}) \in \mathcal{U}^K$, where $\mathcal{U}$ is the finite set of discretized environmental inputs at each step. Its integrated channel capacity is $C_E$, bounded by the physical properties of the respective biological signaling modalities (Berg & Purcell, 1977). We retain the notation $U_{t_0:t_n}$ as a shorthand for the discretized sequence throughout.

4. **Runtime randomness ($W_{0:K-1}$):** Ambient thermal fluctuations, quantum indeterminacy, and uncontrolled molecular collisions acting on the system through its coupling to the physics substrate. Under the time-discretization, $W_{0:K-1} = (W_{\tau_0}, \ldots, W_{\tau_{K-1}}) \in \mathcal{W}^K$ is a finite i.i.d. sequence. From the perspective of the organism's specification scheme, it constitutes an unprogrammed source of bits not contained in $(g, U_{t_0:t_n})$.

5. **Universal compiler and physics substrate:** We distinguish two further objects that are essential to the framework but are *not* signal channels and contribute no bits to the specification budget. The *universal compiler* refers to the laws of physics — electromagnetism, statistical mechanics, chemistry, the structure of the periodic table — which are invariant across all biological systems and which transform low-information specifications into organized microstates. The *physics substrate* refers to the specific physical conditions under which the universal compiler operates for a given biological system: temperature regime, solvent, thermal bath, pressure, density, the existence of the hydrophobic effect, and so on. The laws of physics are universal across all biological systems on Earth, but the substrate is specific to life: living systems occupy a narrow window in the space of physical conditions.

Compilation is conditional on the physics substrate; the universal compiler then ensures that the same specification information evaluated against the same substrate produces a reproducible compilation. Crucially, neither the universal compiler nor the physics substrate carries organism-specific information, neither varies across individuals or developmental contexts, and neither is part of $(C_G + C_E)$. They are the precondition under which the entire framework operates, not contributors to the specification budget. We will return to the substrate distinction when discussing the environmental rescue impossibility (Section "Physical impossibility of the continuous environmental rescue"), where it is essential to be clear that the argument rules out signal-bearing rescue channels, not the role of the universal compiler in transforming specifications into microstate organization.

**Continuous environmental channel and its capacity.** The continuous environmental channel is not meant to denote arbitrary exposure to background physical conditions. It denotes any exogenous, bit-bearing process that crosses the boundary of the biological system and could, in principle, carry organism-specific specifying instructions into the system. This includes morphogens, maternal determinants, hormones, nutrients, pharmacological inputs, bioelectric or mechanical cues, cell–cell signals, and any proposed non-DNA specification substrate, insofar as they function as signals rather than as invariant background physics.

Formally, let $\mathfrak{E}_{\text{bio}}$ be the class of biologically admissible boundary channels over the discretized window $\{\tau_0, \ldots, \tau_{K-1}\}$. A channel $\mathcal{E} \in \mathfrak{E}_{\text{bio}}$ consists of an external source message $M_E$, an encoder producing a boundary sequence $E_{0:K-1} \in \mathcal{V}^K$ (where $\mathcal{V}$ is a finite signal alphabet), a receiver variable $R_{0:K-1}$ inside the system (a function of the trajectory), and a decoder producing $\widehat{M_E}$. Its transmitted specifying information is $I_{\mathcal{E}}(M_E; \widehat{M_E})$, with the supremum taken over admissible message distributions, encoders, receivers, and decoders subject to the physical constraints of the modality: rate, fidelity, addressability, synchronization, and biocompatible actuation. We define

$$C_E := \sup_{\mathcal{E} \in \mathfrak{E}_{\text{bio}}} \sup_{P(M_E),\, \text{enc},\, \text{dec}} I_{\mathcal{E}}(M_E; \widehat{M_E}).$$

This is finite since all variables take values in finite sets. If a modality admits a per-step capacity bound $r_{\mathcal{E}}$ (bits per step), then $C_E \leq K \cdot \sup_{\mathcal{E}} r_{\mathcal{E}}$. In the continuous-time limit, per-step capacity corresponds to an instantaneous rate $r_{\mathcal{E}}(t)$ (bits per second) and the bound becomes $C_E \leq \sup_{\mathcal{E}} \int_{t_0}^{t_n} r_{\mathcal{E}}(t)\, dt$, which is the form used in the modality-specific rate calculations (Supplementary Note 2). In either formulation, $r_{\mathcal{E}}$ is the rate of *addressed and decoded* specifying information, not merely physical energy flux. A broadcast concentration, voltage, strain, light, or temperature field contributes a shared low-dimensional boundary condition unless it can deliver distinct decodable messages to the relevant target cells or subcellular regions.

This definition deliberately excludes the universal compiler and physics substrate. The laws of physics, solvent chemistry, temperature window, hydrophobic effect, and thermal bath are necessary for biological organization, but they are not message channels carrying organism-specific bits. They are part of the fixed background $\Phi$, not part of $C_E$. Likewise, passive environmental variation contributes to $C_E$ only to the extent that it carries mutual information about the target specification. Generic background exposure may alter ensembles, but without decodable specifying information it is not an environmental program.

The physical evolution of this system is governed by two undeniable structural axioms:

**Axiom 1 (Finite state capacity):** Because $\Sigma$ is finite, the maximum forward-control information that can be physically instantiated at any moment $t$ is strictly bounded by $\log_2 |\Sigma|$.

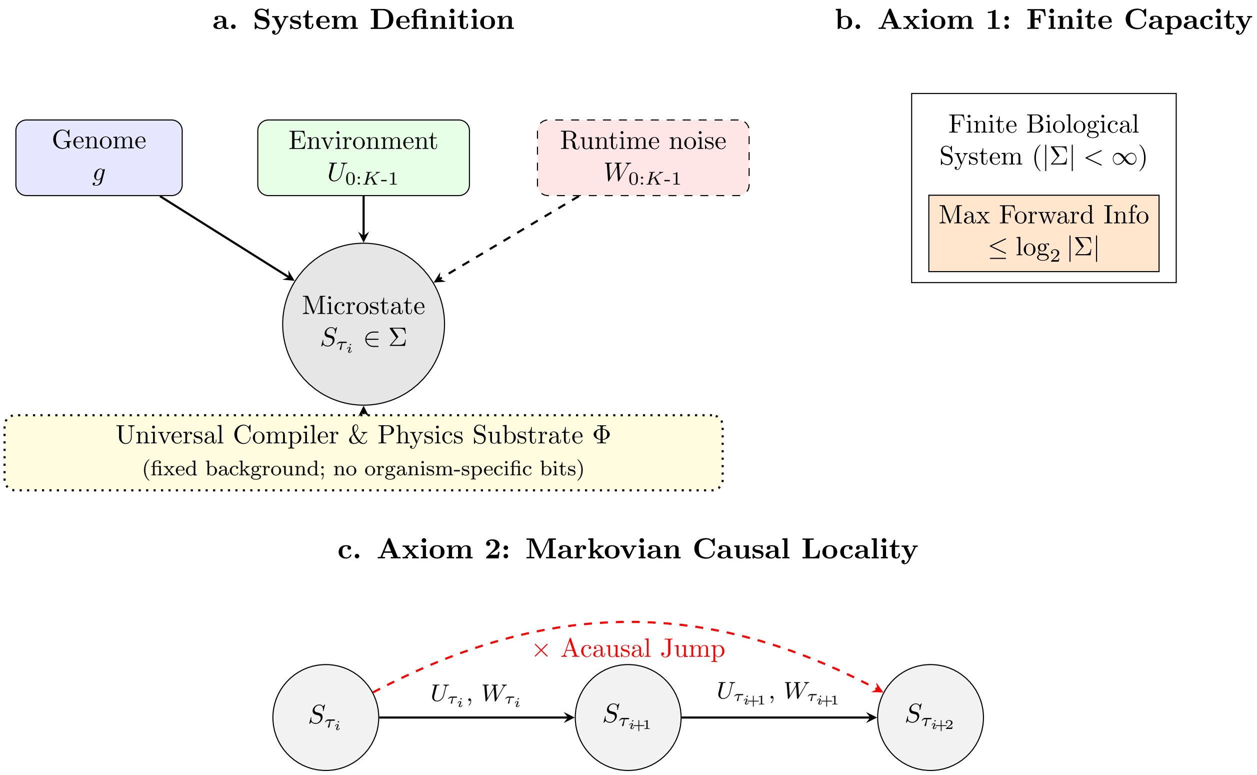


Figure 1: **Formal physical framework of biological specification.** **(a)** The physical microstate trajectory is computed from the static genome $g$, discretized environmental signals $U_{0:K-1}$, thermal runtime randomness $W_{0:K-1}$, and the universal compiler and physics substrate $\Phi$. The genome, environmental channel, and runtime randomness are budgeted or runtime sources; $\Phi$ is held constant as fixed background (dotted border, no organism-specific bits) and is the medium through which the other three operate. **(b)** The physical constraint on forward state capacity. **(c)** Information can only propagate forward locally through intermediate physical states; acausal jumps are forbidden by the Markovian structure of physical dynamics.

**Axiom 2 (Markovian causal locality):** The biological system obeys local physical dynamics. Under the discretization, $S_{\tau_{i+1}}$ is conditionally dependent only on the immediately preceding state $S_{\tau_i}$, the environmental input $U_{\tau_i}$, and the local runtime randomness $W_{\tau_i}$. No forward-directed programmatic instruction can propagate from step $i$ to step $j > i + 1$ without being physically instantiated in all intermediate states.

#### 2 Formal definitions

We now formalize the threshold at which the specification budget is exhausted. All information-theoretic quantities — Shannon entropy $H$, conditional entropy $H(\cdot \mid \cdot)$, mutual information $I$, KL divergence $D$, and Hartley entropy $H_0$ — are defined precisely in Appendix A, together with their application to the specific random variables of the framework. Let $\mathcal{P}$ denote the lattice of all possible coarse-grainings of $\Sigma$.

**Definition 1** (Coarse-graining)**.** A coarse-graining $\mathcal{C} \in \mathcal{P}$ of $\Sigma$ is a partition of $\Sigma$ into disjoint macroscopic cells (i.e., pairwise disjoint subsets; when $\Sigma$ is viewed as the discretization of an underlying compact configuration manifold, these cells correspond to unions of microstate cells at a coarser resolution, equivalently to submanifold-like blocks of the configuration space). A coarse-graining $\mathcal{C}'$ is finer than $\mathcal{C}$ (written $\mathcal{C}' \prec \mathcal{C}$) if every cell of $\mathcal{C}'$ is contained in some cell of $\mathcal{C}$.

**Definition 2** (Induced trajectory and entropy)**.** For a coarse-graining $\mathcal{C}$ of $\Sigma$, the induced discretized trajectory is the random vector $X_{\mathcal{C}} = (\mathcal{C}(S_{\tau_0}), \mathcal{C}(S_{\tau_1}), \ldots, \mathcal{C}(S_{\tau_K})) \in \mathcal{C}^{K+1}$. The entropy of the system at coarse-graining $\mathcal{C}$ over the window is $H(X_{\mathcal{C}})$, which is finite since $X_{\mathcal{C}}$ takes values in a finite set. This entropy is computed under the distribution induced by the physical dynamics (Appendix B). Dependence on the time window $[t_0, t_n]$ and discretization $\delta$ is suppressed in the notation but should be understood throughout.

More generally, a trajectory-level coarse-graining is any partition of the discrete path space $\Sigma^{K+1}$; the pointwise construction above is the special case induced by a partition of $\Sigma$. Since all coarse-grainings used in the empirical sections are of the pointwise form, we work with partitions of $\Sigma$ throughout and note that this covers the relevant cases.

**Remark 1** (Block-spin analogy and monotonicity)**.** The projection $\mathbf{S} \mapsto X_{\mathcal{C}}$ is formally analogous to the block-spin (real-space renormalization group) operation of statistical physics: it is a many-to-one map on discrete path space $\Sigma^{K+1}$, and is therefore non-invertible. Passing from a finer coarse-graining $\mathcal{C}'$ to a coarser one $\mathcal{C}$ (i.e., along $\mathcal{C}' \prec \mathcal{C}$) is well-defined and entropy-monotone, $H(X_{\mathcal{C}'}) \geq H(X_{\mathcal{C}})$; passing from coarser to finer is not, since fine-grained paths are not recoverable from their coarse-grained projections. The lattice $\mathcal{P}$ of coarse-grainings inherits this one-way structure.

**Proposition 1** (Monotonicity under refinement)**.** *If $\mathcal{C}' \prec \mathcal{C}$, let*

$$q_{\mathcal{C}\leftarrow\mathcal{C}'} : \mathcal{C}' \longrightarrow \mathcal{C}$$

*be the surjection sending each fine cell to the unique coarse cell containing it. Then, denoting by $\pi_{\mathcal{C}\leftarrow\mathcal{C}'} : (\mathcal{C}')^{K+1} \to \mathcal{C}^{K+1}$ the resulting map on trajectories,*

$$X_{\mathcal{C}} = \pi_{\mathcal{C}\leftarrow\mathcal{C}'} \circ X_{\mathcal{C}'} \quad \textit{almost surely,}$$

*and consequently*

$$H(X_{\mathcal{C}}) \leq H(X_{\mathcal{C}'}).$$

*Proof.* Because every cell of the finer partition $\mathcal{C}'$ is contained in exactly one cell of the coarser partition $\mathcal{C}$, each $\mathcal{C}'$-cell determines a unique $\mathcal{C}$-cell. This induces a deterministic surjection from $\mathcal{C}'$-symbols to $\mathcal{C}$-symbols, and therefore a deterministic map on trajectory-valued random variables,

$$X_{\mathcal{C}} = \pi_{\mathcal{C}\leftarrow\mathcal{C}'}(X_{\mathcal{C}'}).$$

Since deterministic maps cannot increase Shannon entropy, the data-processing inequality yields

$$H(X_{\mathcal{C}}) \leq H(X_{\mathcal{C}'}).$$

□

**Definition 3** (Programmed microstate determinism)**.** A biological system exhibits programmed microstate determinism at coarse-graining $\mathcal{C}$ if its realized trajectory $X_{\mathcal{C}}$ is determined by the joint specification $(g, U_{t_0:t_n})$. This admits two precise formulations:

- *Hartley form:* for every admissible specification $(g, u)$, the conditional support is a singleton, $|\operatorname{supp}(X_{\mathcal{C}} \mid g = g, U = u, \Phi)| = 1$. That is, the specification pins down a unique trajectory.
- *Shannon form:* the conditional Shannon entropy vanishes, $H(X_{\mathcal{C}} \mid g, U_{t_0:t_n}) = 0$, under the probability model of Appendix B. That is, $X_{\mathcal{C}}$ is almost surely a deterministic function of $(g, U_{t_0:t_n})$.

The Hartley form implies the Shannon form. Both are properties of being specifiable by the organism's own information stores. Both are distinct from physical determinism, which concerns whether the system's microstate evolves deterministically under the laws of physics independent of any specification scheme. The theorem of this paper addresses programmed microstate determinism only.

### 3 Core lemmas

**Lemma 1 (Runtime-randomness lower bound).** *Let $A$ be any algorithm that, on input $(g, U_{t0:t_n})$, reads i.i.d. fair random bits until halting and outputs $X \in \Sigma$. Let $T$ denote the number of random bits consumed. Then for every fixed $(g, U_{t0:t_n})$,*

$$\mathbb{E}[T \mid g, U_{t0:t_n}] \geq H(X \mid g, U_{t0:t_n}).$$

*Proof.* Let $\mathcal{L} \subseteq \{0,1\}^*$ be the set of halting random-bit transcripts. Because the algorithm halts after reading a finite prefix, $\mathcal{L}$ is prefix-free. Let $Y \in \mathcal{L}$ be the realized transcript. Then $\Pr(Y = y) = 2^{-|y|}$ for each $y \in \mathcal{L}$, so $H(Y) = \sum_{y \in \mathcal{L}} 2^{-|y|}|y| = \mathbb{E}|Y| = \mathbb{E}[T]$. Since $X$ is a deterministic function of $Y$ once $(g, U_{t0:t_n})$ are fixed, the data-processing inequality gives $H(X \mid g, U_{t0:t_n}) \leq H(Y \mid g, U_{t0:t_n}) = \mathbb{E}[T \mid g, U_{t0:t_n}]$. □

A skeptic may object that the deterministic program is highly compressed and that the genome *is* the compressed form. The lemma closes this loophole: a compressed program with a stochastic decompressor must consume runtime randomness at least equal to the output entropy. Bits can be relocated from explicit description into runtime randomness, but they cannot be made to vanish. The lemma is also agnostic about the ultimate origin of the runtime random bits, which may come from genuine quantum indeterminacy, thermal fluctuations that are physically deterministic but uncontrolled, or any other source the organism does not specify. What matters is that they are not in $(g, U_{t0:t_n})$.

**Connection to the physical dynamics.** Under the time-discretization of Section "Formal physical framework", each transition $S_{\tau_{i+1}} \mid (S_{\tau_i}, U_{\tau_i})$ is a random variable on $\Sigma$ whose conditional law is determined by the Markov kernel. Sampling from this conditional law requires at most $\log_2 |\Sigma|$ fair bits per step (via the inverse-CDF construction; see Appendix C). Over $K$ steps, the total randomness consumed is $T \leq K \log_2 |\Sigma|$, and Lemma 1 applies with $X = \mathbf{S} = (S_{\tau_0}, \ldots, S_{\tau_K})$. The thermal noise $W_t$ of the physical system plays the role of the i.i.d. fair bits in the lemma: it is the source from which each transition draws the randomness not determined by $(g, U)$.

**Initial conditions and the decay of predictive information.** The runtime-randomness lemma bounds the random bits consumed given $(g, U_{t0:t_n})$. A deeper objection is that the full initial microstate $S_{\tau_0}$ contains up to $\log_2 |\Sigma|$ bits, and this large static information store could in principle seed a deterministic trajectory without consuming fresh runtime random bits. We close this loophole by proving that the predictive information carried by $S_{\tau_0}$ decays exponentially under the driven thermal dynamics.

Assume uniform KL contraction for the discretized driven chain: for all steps $j \geq i$ and all probability measures $\mu, \nu$ on $\Sigma$, the multi-step Markov kernel $K_{i,j}^{g,U}$ (the composition of single-step kernels from step $i$ to step $j$) satisfies

$$D(\mu K_{i,j}^{g,U} \| \nu K_{i,j}^{g,U}) \leq e^{-(j-i)\delta/\tau_{\text{mix}}} D(\mu \| \nu),$$

where $D(\cdot\|\cdot)$ denotes the Kullback–Leibler divergence, $\delta$ is the time-discretization step, and $\tau_{\text{mix}} > 0$ is the contraction timescale (in physical time units). This is a uniform (all-pairs) KL contraction hypothesis, stronger than contraction relative to the stationary measure alone; for finite irreducible chains it is equivalent to uniform ergodicity. The "uniform" qualifier does genuine work: the bound applies to all pairs $(\mu, \nu)$, not only to $(\mu, \pi)$ where $\pi$ is the stationary distribution. For a time-homogeneous chain, the contraction rate is controlled by the spectral gap of the generator; for the driven chain (where transition rates depend on the time-varying $U_{\tau_i}$), the hypothesis requires that the environmental signal does not break ergodicity. This is physically reasonable for biological systems in or near thermal equilibrium under slowly varying environmental conditions (see Appendix E).

**Lemma 1b (Initial-condition mixing).** *Under the uniform KL contraction assumption, the predictive mutual information carried by the initial microstate decays exponentially: for any step* $j \geq 0$,

$$I(S_{\tau_0}; S_{\tau_j} \mid g, U_{0:j-1}) \leq e^{-j\delta/\tau_{\text{mix}}} \log_2 |\Sigma|.$$

*Proof.* Fix $g$ and $U_{0:j-1}$, let $\mu_0$ denote the conditional law of $S_{\tau_0}$, and write $K := K^{g,U}_{0,j}$. Then $I(S_{\tau_0}; S_{\tau_j} \mid g, U_{0:j-1}) = \sum_x \mu_0(x) D(\delta_x K \| \mu_0 K)$. Applying the contraction hypothesis with $\mu = \delta_x$ and $\nu = \mu_0$ gives $D(\delta_x K \| \mu_0 K) \leq e^{-j\delta/\tau_{\text{mix}}} D(\delta_x \| \mu_0)$. Averaging over $x \sim \mu_0$ yields $I(S_{\tau_0}; S_{\tau_j} \mid g, U_{0:j-1}) \leq e^{-j\delta/\tau_{\text{mix}}} H(S_{\tau_0} \mid g, U_{0:j-1}) \leq e^{-j\delta/\tau_{\text{mix}}} \log_2 |\Sigma|$. □

**Corollary (Runtime-randomness with initial-state side information).** *For any output $X$ generated after $j$ steps (physical burn-in time $j\delta$), $\mathbb{E}[T \mid g, U, S_{\tau_0}] \geq H(X \mid g, U) - e^{-j\delta/\tau_{\text{mix}}} \log_2 |\Sigma|$. In particular, if $e^{-j\delta/\tau_{\text{mix}}} \log_2 |\Sigma| \leq \varepsilon$, the original runtime-randomness bound holds up to an additive $\varepsilon$-bit correction.*

For diffusive microscopic degrees of freedom associated with molecular length scale $a$, the Stokes-Einstein relation gives a single-particle positional relaxation time $\tau_{\text{SE}} \sim 6\pi\eta a^3/(k_B T)$, i.e., the time for a molecule of radius $a$ to diffuse across a distance of order $a$. At $T = 310$ K with water-like viscosity, this yields $\tau_{\text{SE}} \sim 10^{-10}$ to $10^{-8}$ s for molecular scales $a \in [0.3, 1]$ nm. For independent particles, the full-state contraction timescale $\tau_{\text{mix}}$ equals the single-particle timescale (independent processes mix in parallel). Inter-molecular interactions can create slower collective modes; the slow/fast decomposition below handles this by restricting the contraction hypothesis to the fast residual degrees of freedom. Cytoplasmic viscosity is not water viscosity: small-probe measurements give effective viscosities of roughly 2–10× water in bulk cytoplasm, with values reaching up to $\sim 10^3\times$ water in crowded or gel-like subcellular regions. Even under the most adverse estimate (inflating $\tau_{\text{mix}}$ by three orders of magnitude to $\sim 10^{-5}$ s), the exponent $j\delta/\tau_{\text{mix}}$ exceeds $10^3$ for any $j\delta$ on the millisecond scale, making $e^{-j\delta/\tau_{\text{mix}}}$ astronomically small. Any millisecond-to-hour biological window lies overwhelmingly in the regime $j\delta \gg \tau_{\text{mix}}$. The initial-condition loophole is closed for these fast residual modes.

**Remark 2** (Scope of the mixing lemma)**.** Lemma 1b is intended to close the Laplacian-demon loophole for *fast* residual degrees of freedom whose predictive information would otherwise be attributed to the initial condition. Long-lived structured memory — for example chromatin organization, epigenetic marks, slowly varying condensates, membrane domains, cytoskeletal assemblies, or other actively maintained slow manifolds — is not claimed to mix on nanosecond scales. Such slow variables are part of the current physical state itself and are therefore governed by Lemma 2 rather than by Lemma 1b: they count as state-carried memory competing with current function for the same finite causal-locality budget. Thus slow biological memory does not rescue a hidden initial-condition program; it is already one of the finite stores whose capacity the framework bounds.

**Scoped form of the mixing lemma.** The verbal scope above admits a precise formal statement via a slow/fast decomposition of the microstate. Write

$$S_{\tau_i} = (Y_{\tau_i}, Z_{\tau_i}),$$

where $Y_{\tau_i}$ denotes slow, state-carried memory variables whose autocorrelation time exceeds a biological relevance threshold $\tau_{\text{bio}}$ (chromatin, epigenetic marks, slow condensates, cytoskeletal assemblies), and $Z_{\tau_i}$ denotes fast residual variables. We assume uniform KL contraction *only* for the conditional fast dynamics,

$$D\big(\mu K_{i,j}^{g,U,Y} \,\big\|\, \nu K_{i,j}^{g,U,Y}\big) \;\le\; \exp[-(j-i)\delta/\tau_{\text{fast}}]\, D(\mu\|\nu),$$

where $K_{i,j}^{g,U,Y}$ is the multi-step Markov kernel for $Z$ conditional on the slow path $Y_{\tau_i:\tau_j}$. Then

$$I(S_{\tau_0}; S_{\tau_j} \mid g, U_{0:j-1}) \;\le\; H(Y_{\tau_0} \mid g, U_{0:j-1}) + \exp[-j\delta/\tau_{\text{fast}}] \log_2 |\Sigma_Z|.$$

The first term is slow state-carried memory, governed by causal locality (Lemma 2) rather than by mixing. The second term is the fast initial-condition residue, which decays exponentially. Slow memory is therefore not a loophole: it is precisely the part of the current state that Lemma 2 already counts as finite state-carried information, and it remains subject to that bound.

**Lemma 2 (Causal locality).** *Let $(S_k)_{k=0}^n$ be the discretized trajectory on $\Sigma$, with dynamics Markov in $S_k$: conditional on $(S_{k-1}, U_k, W_k)$, the state $S_k$ is independent of $(S_0, \ldots, S_{k-2}, g)$. Let $\phi : \Sigma \times \mathcal{U}^{n-k+1} \to \mathcal{A}$ be any measurable function (representing a deterministic forward program that depends on the current state and the future environmental signal but not on future noise). Define $A_{k:n} = \phi(S_{k-1}, U_{k:n})$. Then*

$$H(A_{k:n} \mid U_{k:n}) \le H(S_{k-1} \mid U_{k:n}) \le \log_2 |\Sigma|.$$

*More generally, for any pair of measurable functions $f : \Sigma \to \mathcal{F}$ ("current function") and $\phi : \Sigma \times \mathcal{U}^{n-k+1} \to \mathcal{A}$ ("forward program"),*

$$H(f(S_{k-1}), \phi(S_{k-1}, U_{k:n}) \mid U_{k:n}) \le H(S_{k-1} \mid U_{k:n}) \le \log_2 |\Sigma|.$$

*Proof.* Since $A_{k:n} = \phi(S_{k-1}, U_{k:n})$ is a deterministic function of $(S_{k-1}, U_{k:n})$, the data-processing inequality gives $H(A_{k:n} \mid U_{k:n}) \le H(S_{k-1}, U_{k:n} \mid U_{k:n}) = H(S_{k-1} \mid U_{k:n})$. The second inequality follows from $H(S_{k-1} \mid U_{k:n}) \le H(S_{k-1}) \le \log_2 |\Sigma|$. The general statement follows identically, since the pair $(f(S_{k-1}), \phi(S_{k-1}, U_{k:n}))$ is also a deterministic function of $(S_{k-1}, U_{k:n})$. □

The key content of the lemma is the definition of "programmed": a forward action $A_{k:n}$ is *programmed* if and only if it is a deterministic function of the current state $S_{k-1}$ and future environmental inputs $U_{k:n}$, with no dependence on future thermal noise $W_{k:n}$. Any information that a programmed action requires must therefore be physically present in $S_{k-1}$ at the moment the action is initiated, where it competes with current physiological function for the same finite capacity $\log_2 |\Sigma|$. The full conditional-independence structure underlying this statement is recorded in Appendix F.

The lemma is fully compatible with biological state-carried memory: phosphorylation, epigenetic marks, synaptic weights, and membrane potential are all part of $S_t$ and are counted by the same bound. What it forbids is delayed instructions whose execution at $t_k$ is indexed to a future wall-clock time without an intervening physical trigger in the state at $t_{k-1}$. This also resolves the DNA-as-blueprint objection: DNA is compatible with the lemma as a *generator specification* (a gene sequence specifies a molecular machine $M$ whose local kinetics produce behavior $B$ on the current substrate), but not as a free delayed-trajectory tape, since transcription is initiated by local promoter recognition, RNA polymerase availability, and chromatin accessibility (Munsky *et al.*, 2012), all of which are state variables, rather than by a global clock indexing into the genome sequence.

### 4 Zero-error addressability and the cost of reducing a fine-state fiber

The two foregoing lemmas constrain randomness consumption (Lemma 1, Lemma 1b) and forward-control capacity (Lemma 2). A third constraint, complementary to both, governs the *deterministic addressability* of fine states: how many bits an organism-controlled specification would have to expend to select an arbitrary distinguishable fine-grained realization within the fiber left unresolved by a functional coarse-graining. This is a zero-error coding statement, distinct from Shannon entropy, and it is the appropriate object whenever the empirical claim concerns deterministic specification rather than stochastic sampling.

**Definition 4** (Admissible fiber and zero-error support cost)**.** Fix the organism-controlled specification together with the shared physical background, $z = (g, U_{t_0:t_n}, \Phi)$, where $\Phi$ denotes the universal compiler and physics substrate as defined above. For a coarse-graining $\mathcal{C}$, define the admissible fine-state fiber

$$\mathcal{A}_\mathcal{C}(z) \;=\; \operatorname{supp}\big(X_\mathcal{C} \,\big|\, z\big),$$

i.e. the set of fine-grained realizations of $X_\mathcal{C}$ that remain consistent with $z$. The zero-error support cost (Hartley entropy conditioned on $z$) is

$$H_0\big(X_\mathcal{C} \,\big|\, z\big) \;=\; \log_2\big|\mathcal{A}_\mathcal{C}(z)\big|.$$

This is the number of bits required to address an arbitrary distinguishable fine-grained realization in the fiber. It is distinct from the Shannon conditional entropy $H(X_\mathcal{C} \mid z)$, which is the expected number of random bits required to sample from the conditional distribution on the fiber, and the two satisfy $H(X_\mathcal{C} \mid z) \le H_0(X_\mathcal{C} \mid z)$ in general, with equality only when the conditional distribution is uniform on $\mathcal{A}_\mathcal{C}(z)$.

**Proposition 2** (Zero-error addressability bound)**.** *Fix $z = (g, U_{t_0:t_n}, \Phi)$. Suppose a deterministic specifier uses a message $M$ of length at most $L$ bits, together with side information $R$ having conditional support of size $|\mathcal{R}_z|$, to select states in $\mathcal{A}_\mathcal{C}(z)$ with zero error. Then*

$$L + H_0\big(R \,\big|\, z\big) \;\ge\; H_0\big(X_\mathcal{C} \,\big|\, z\big).$$

*In particular, when no side information is available, exact deterministic addressability of every state in $\mathcal{A}_\mathcal{C}(z)$ requires at least* $\log_2 |\mathcal{A}_\mathcal{C}(z)|$ *bits.*

*Proof.* For fixed $z$, a message of length at most $L$ takes at most $2^L$ possible values. If the side information $R$ has conditional support $\mathcal{R}_z$, then the pair $(M, R)$ takes at most $2^L|\mathcal{R}_z|$ possible values. A deterministic decoder maps each pair $(m, r)$ to at most one fine state. Therefore it can select at most $2^L|\mathcal{R}_z|$ distinct fine states. To address every state in $\mathcal{A}_\mathcal{C}(z)$, one must have $|\mathcal{A}_\mathcal{C}(z)| \le 2^L|\mathcal{R}_z|$. Taking $\log_2$ gives the stated inequality. □

**Corollary 1** (Cost of reducing a fine-state fiber). *Let $R_t \subseteq S_t$ be a state-carried biological variable that reduces the admissible fiber from $\mathcal{A}_{\mathcal{C}}(z)$ to conditional fibers $\mathcal{A}_{\mathcal{C}}(z, r)$. If $\max_r \log_2 |\mathcal{A}_{\mathcal{C}}(z, r)| \leq h$, then*

$$H_0\big(R_t \,\big|\, z\big) \;\geq\; H_0\big(X_{\mathcal{C}} \,\big|\, z\big) - h.$$

*In particular, exact selection of a single fine state ($h = 0$) requires $H_0(R_t \mid z) \geq H_0(X_{\mathcal{C}} \mid z)$. Any biological mechanism that reduces the unresolved fine-state support by $\beta$ bits must therefore physically carry those $\beta$ bits in $g$, $U$, or the current state $S_t$, where they are counted by the specification budget and by causal locality.*

*Proof.* The same covering argument: if each $r \in \mathcal{R}_z$ leaves at most $2^h$ admissible fine states, and $R_t$ takes at most $2^{H_0(R_t|z)}$ values, then the union of $r$-conditional fibers covers at most $2^{H_0(R_t|z)+h}$ fine states. To cover $\mathcal{A}_{\mathcal{C}}(z)$, $H_0(R_t \mid z) + h \geq H_0(X_{\mathcal{C}} \mid z)$. □

The corollary identifies precisely where any putative "fiber-reducing" mechanism must reside: in $g$, in $U$, in the current state $S_t$ (subject to Lemma 2), in the universal compiler $\Phi$ (which is shared and not organism-specific), or in runtime randomness (which is not programmed). There is no fourth category. This is the formal foreclosure of the natural objection that folded proteins, scaffold assemblies, or other state-carried structures somehow reduce the spatial-organization fiber "for free": they do reduce it, but the reduction is paid for in side-information bits subject to the same lemmas that govern all forward control.

The relationship between the two information quantities is given by

$$H_0\big(X \,\big|\, z\big) - H\big(X \,\big|\, z\big) \;=\; D\big(P_{X|z} \,\big\|\, U_{\mathcal{A}(z)}\big)\,,$$

where $U_{\mathcal{A}(z)}$ is the uniform distribution on $\mathcal{A}(z)$ and $D$ is the Kullback–Leibler divergence. The gap between zero-error support cost and Shannon entropy is therefore exactly the amount of distributional structure in the conditional law on the fiber. Any such structure is itself information, classified by the same dichotomy: supplied by the specification budget $C_G + C_E$, by state-carried memory (Lemma 2), by the shared compiler $\Phi$ (which is not organism-specific), or by runtime fluctuations (which are not programmed).

The combinatorial calculations in the empirical sections below should be read as zero-error addressability bounds on $H_0(X_{\mathcal{C}} \mid z)$, not as lower bounds on Shannon entropy of the realized distribution. Any mechanism that reduces a fine-state fiber must carry the corresponding side information in the genome, the environmental signal, or the current physical state, where it is counted by finite capacity and causal locality; otherwise the reduction is supplied by the shared compiler or runtime physics and is not organism-programmed. The Shannon-entropy version of the impossibility statement is supplied separately by the runtime-randomness lemma, and the two impossibility statements together — zero-error addressability and runtime-randomness cost — form the two prongs of the coarse-graining theorem below.

### 5 The coarse-graining threshold

We have established two complementary information-theoretic constraints on organism-controlled specification: the runtime-randomness lemma (Lemma 1, Lemma 1b), which bounds the random bits required to sample from a conditional distribution, and the zero-error addressability proposition, which bounds the bits required to select an arbitrary distinguishable fine state. Both speak to the same underlying question — what can the organism program at a given coarse-graining?

— but they answer it through different information-theoretic objects, and they apply to different specification tasks. We now combine them into a single threshold concept and then specialize it.

**Notation: budgeted message and fixed background.** Throughout this subsection we separate the organism-controlled budgeted message

$$M = (g, U_{t_0:t_n}),$$

from the fixed, non-message background

$$b \; = \; (\Phi, \text{system identity}, \text{observation window } [t_0, t_n], \text{discretization } \delta),$$

which is held constant in the comparison and which contributes no organism-specific bits. The total specification budget is $B = C_G + C_E$. For the Hartley prong, $B$ bounds the log-cardinality of the message space: a genome of $n$ bp admits at most $4^n$ values, and the environmental channel admits at most $2^{C_E}$ distinguishable signals, so $|\operatorname{supp}(M)| \leq 2^B$. For the Shannon prong, $B$ upper-bounds the Shannon entropy $H(M)$ (since $H(M) \leq \log_2 |\operatorname{supp}(M)| \leq B$; see Appendix D). This separation makes the threshold inequalities transparent: thresholds compare the pre-message information cost of $X_{\mathcal{C}}$ given only the fixed background $b$ to the message budget $B$, without conditioning on $M$ inside the threshold itself.

**The unified threshold.** Let $\mathcal{H}(X_{\mathcal{C}} \mid b)$ denote a generic information functional measuring the number of bits required to specify $X_{\mathcal{C}}$ given the fixed background $b$. Define the set of capacity-compatible coarse-grainings as

$$\mathfrak{C}_{\mathcal{H}}(B) \; = \; \{\mathcal{C} \in \mathcal{P} : \mathcal{H}(X_{\mathcal{C}} \mid b) \leq B\},$$

and let $\mathfrak{C}^*_{\mathcal{H}}(B)$ be the antichain of maximal elements of $\mathfrak{C}_{\mathcal{H}}(B)$ under refinement. The threshold is the boundary of this antichain: above it, organism-controlled specification at resolution $\mathcal{C}$ is not ruled out by capacity alone; below it, the specification task requires more bits than the message budget $B$ contains.

This single threshold concept admits two natural formal expressions, depending on whether the specification task is stochastic sampling or deterministic addressability. They correspond to two standard information entropies:

- $\mathcal{H} = H$, the *Shannon* conditional entropy $H(X_{\mathcal{C}} \mid b)$, governing the average runtime randomness required to sample from the realized distribution. This is the appropriate entropy when the specification task is stochastic ensemble generation: how many random bits must be consumed beyond what the message $M$ carries.
- $\mathcal{H} = H_0$, the *Hartley* zero-error support cost $H_0(X_{\mathcal{C}} \mid b) = \log_2 |\operatorname{supp}(X_{\mathcal{C}} \mid b)|$, governing the bits required to address an arbitrary distinguishable fine state. This is the appropriate entropy when the specification task is deterministic selection: whether the organism can pick out one fine realization from the admissible set.

These two specializations give two threshold families, both special cases of the unified definition above:

$$\mathfrak{C}_1(B) \; = \; \{\mathcal{C} \in \mathcal{P} : H(X_{\mathcal{C}} \mid b) \leq B\},$$

$$\mathfrak{C}_0(B) \; = \; \{\mathcal{C} \in \mathcal{P} : H_0(X_{\mathcal{C}} \mid b) \leq B\}.$$

The family $\mathfrak{C}_1$ is the Shannon-compatible family, relevant for stochastic generation; $\mathfrak{C}_0$ is the Hartley-compatible family, relevant for zero-error deterministic addressability. Since $H(X_\mathcal{C} \mid b) \leq H_0(X_\mathcal{C} \mid b)$ for any random variable on a finite support, we have the nesting

$$\mathfrak{C}_0(B) \subseteq \mathfrak{C}_1(B),$$

i.e. zero-error compatibility implies Shannon compatibility, and the zero-error threshold is at least as coarse as the Shannon threshold. Let $\mathfrak{C}_1^*(B)$ and $\mathfrak{C}_0^*(B)$ denote the corresponding antichains of maximal elements under refinement. Between the two thresholds lies a biologically meaningful middle zone where stochastic ensemble generation can fit within the message budget but deterministic zero-error addressability cannot.

The threshold is therefore a single conceptual object with two standard information-theoretic faces: a Shannon face, controlling stochastic generation, and a Hartley face, controlling zero-error deterministic addressability. The former asks how many random bits are needed to sample the ensemble; the latter asks how many bits are needed to select an exact fine state. Biology, as the empirical sections will show, lives above both thresholds at the functional level and below both at microstate resolution.

**Theorem 1 (Coarse-graining threshold for programmed specification).** *Let $\mathcal{P}$ be the lattice of coarse-grainings of $\Sigma$ ordered by refinement, let $b$ denote the fixed non-message background, let $M = (g, U_{t_0:t_n})$ denote the budgeted organism-controlled message with $|\operatorname{supp}(M)| \leq 2^B$ where $B = C_G + C_E$. Then:*

1. *(Capacity-compatible regime.) If $\mathcal{C} \in \mathfrak{C}_1(B)$, then stochastic specification at resolution $\mathcal{C}$ is not ruled out by Shannon capacity alone: there exists an abstract lossless source code for $X_\mathcal{C}$ whose expected code length is at most $H(X_\mathcal{C} \mid b) + 1 \leq B + 1$ bits. If $\mathcal{C} \in \mathfrak{C}_0(B)$, then zero-error deterministic addressability is not ruled out by Hartley capacity alone. Neither condition is sufficient for biological realization; sufficiency additionally requires that the corresponding code is present in $M$, causally exposed through state, and physically realizable.*
2. *(Sub-threshold impossibility, two prongs.) Let $\mathcal{C}'$ be a strict refinement below the relevant threshold antichain.*
   (a) *Zero-error addressability: if $\mathcal{C}'$ is a strict refinement of some $\mathcal{C}_0^* \in \mathfrak{C}_0^*(B)$ and $H_0(X_{\mathcal{C}'} \mid b) > B$, then no organism-controlled deterministic specification of capacity $B$ can address every distinguishable fine state in $\operatorname{supp}(X_{\mathcal{C}'} \mid b)$.*
   (b) *Runtime-randomness cost: if $\mathcal{C}'$ is a strict refinement of some $\mathcal{C}_1^* \in \mathfrak{C}_1^*(B)$ and $H(X_{\mathcal{C}'} \mid b) > B$, then $H(X_{\mathcal{C}'} \mid M, b) \geq H(X_{\mathcal{C}'} \mid b) - B$, and any algorithm producing $X_{\mathcal{C}'}$ from $M$ must consume runtime randomness with expected length at least $H(X_{\mathcal{C}'} \mid b) - B$.*

*Proof.* For (1): the first half follows from the noiseless source-coding theorem (Shannon, 1948; Cover & Thomas, 2006), applied to $X_\mathcal{C}$ given $b$: there exists a prefix code of expected length strictly less than $H(X_\mathcal{C} \mid b) + 1$. Thus, when $H(X_\mathcal{C} \mid b) \leq B$, an abstract one-shot prefix description lies within one bit of the budget; under block coding, the per-sample rate approaches $H(X_\mathcal{C} \mid b) \leq B$. The second half follows from the zero-error addressability bound: if $H_0(X_\mathcal{C} \mid b) \leq B$, then $|\operatorname{supp}(X_\mathcal{C} \mid b)| \leq 2^B$, and a deterministic specifier with message of length $B$ can in principle address every element. Sufficiency in either case additionally requires (i) that the required mutual information $I(X_\mathcal{C}; M \mid b)$ actually equals $\mathcal{H}(X_\mathcal{C} \mid b)$, (ii) that the code is causally realizable, and (iii) that the resulting decoder is physically realizable under the Markov dynamics on $\Sigma$.

For (2a): by the zero-error addressability bound (Proposition 2), any deterministic specifier using a message $M$ of length at most $B$ bits can select at most $2^B$ distinct fine states. If $H_0(X_{\mathcal{C}'} \mid b) > B$, then $|\operatorname{supp}(X_{\mathcal{C}'} \mid b)| > 2^B$, so no such specifier can address every state in the admissible fiber. By the side-information corollary, any putative fiber-reducing variable $R_t \subseteq S_t$ must itself carry $H_0(R_t \mid b) \geq H_0(X_{\mathcal{C}'} \mid b) - h$ bits, which by Lemma 2 are bounded by $\log_2 |\Sigma|$ and compete with current function for the same instantaneous capacity. Therefore no organism-controlled deterministic specification at message capacity $B$ exhausts the fiber.

For (2b): since $|\operatorname{supp}(M)| \leq 2^B$, we have $H(M) \leq \log_2 |\operatorname{supp}(M)| \leq B$ under any distribution on $M$ (Appendix D). Therefore $I(X_{\mathcal{C}'}; M \mid b) \leq H(M) \leq B$, so $H(X_{\mathcal{C}'} \mid M, b) = H(X_{\mathcal{C}'} \mid b) - I(X_{\mathcal{C}'}; M \mid b) \geq H(X_{\mathcal{C}'} \mid b) - B > 0$. The runtime-randomness lemma (Lemma 1) then gives $\mathbb{E}[T \mid M, b] \geq H(X_{\mathcal{C}'} \mid M, b) \geq H(X_{\mathcal{C}'} \mid b) - B$. (For a fixed genome $g$, the bound simplifies to $H(X_{\mathcal{C}'} \mid g, U, b) \geq H(X_{\mathcal{C}'} \mid g, b) - C_E$, where only the environmental channel capacity appears.) □

The two prongs of Part (2) answer two distinct questions through their respective entropies. Prong (2a) asks whether the organism can deterministically address an arbitrary fine state in the unresolved fiber, and uses Hartley entropy. Prong (2b) asks how much runtime randomness must be consumed if the organism instead samples from a particular conditional distribution, and uses Shannon entropy. Both prongs are special cases of the unified threshold concept and use information-theoretic objects appropriate to the question they answer; neither relies on the empirical conditional distribution being uniform on its support.

**Approximate addressability.** Both threshold faces admit a natural relaxation to the case where the specifier tolerates a small error probability. By Fano's inequality, any function selecting one of $K$ equiprobable fine states with error probability $\varepsilon$ requires mutual information at least $(1 - \varepsilon) \log_2 K - h_2(\varepsilon)$ bits, where $h_2$ is the binary entropy function. Allowing a small error probability therefore softens the zero-error bound by $O(h_2(\varepsilon) + \varepsilon \log K)$, but does not change the order-of-magnitude conclusion for the biological gaps considered here, where the support cardinalities exceed organism-controlled capacity by factors of 10 or more. The framework's impossibility statements remain intact under approximate addressability with realistic biological error tolerances.

**What a sufficiency proof would require.** The gap between Part (1) (capacity-compatibility) and full programmed determinism ($H(X_{\mathcal{C}} \mid M, b) = 0$) is closed by constructing a *realizable causal code*: a message $M$ with $H(M) \leq B$ and $H(X_{\mathcal{C}} \mid M, b) = 0$, a causal exposure rule ensuring $M$ can be read from $S_{\tau_i}$ at each step without depending on future inputs, and a physical realizability condition ensuring the Markov dynamics on $\Sigma$ can implement the decoding. For static or endpoint coarse-grainings (where no intermediate forward schedule must be carried through the state history), the existence of a deterministic readout suffices and the causal-locality constraint imposes no additional runtime schedule; this restricted sufficiency holds whenever $H(X_{\mathcal{C}} \mid b) \leq B$. For trajectory-level coarse-grainings, the construction is genuinely harder and is not provided here. The empirical content of the paper does not depend on the sufficiency direction: the central claim is that sub-threshold specification is impossible (Part 2), not that above-threshold specification is guaranteed.

The threshold is in general not a single canonical coarse-graining but an antichain of maximal compatible coarse-grainings. There are typically multiple mutually incomparable ways to maximally fill the budget — one element of $\mathfrak{C}_0^*(B)$ may permit deterministic addressability of certain fine states while another permits deterministic addressability of different fine states, with similar incomparability among elements of $\mathfrak{C}_1^*(B)$. The empirical claim of the next sections is that biological function is defined at coarse-grainings that lie within these antichains.

**Notation for empirical comparisons.** In all numerical comparisons in the empirical sections, we use the raw haploid genome bit count

$$C_G^{\mathrm{raw,hap}} \;=\; 2\cdot(\text{haploid bp})$$

as a generous upper bound on $C_G$, where the factor of two reflects two bits per nucleotide ($\log_2 4$ for the four-letter DNA alphabet). The information content of the heritable store is bounded above by this quantity regardless of ploidy. To see this for sexually reproducing diploid organisms, let $g_1, g_2$ denote the two homologous genome copies and let $\eta$ denote the within-individual heterozygosity rate (the fraction of positions at which the two homologs differ). Then by the chain rule of entropy and the fact that conditioning reduces entropy,

$$C_G^{\mathrm{raw,dip}} \;=\; H(g_1, g_2) \;=\; H(g_1) + H(g_2 \mid g_1) \;\le\; (1+\eta)\, C_G^{\mathrm{raw,hap}},$$

since the bits required to specify $g_2$ given $g_1$ are at most the bits required to specify the differing positions. With $\eta < 0.01$ in humans (1000 Genomes Project Consortium, 2015) and lower in many other species, $C_G^{\mathrm{raw,dip}} \lesssim 1.01\, C_G^{\mathrm{raw,hap}}$, so the haploid bound is also a tight upper bound on the diploid information content, to within at most 1%. Combining with the obvious bound that functional capacity cannot exceed raw bit count,

$$C_G^{\mathrm{func}} \;\le\; C_G^{\mathrm{raw,hap}} \;\lesssim\; C_G^{\mathrm{raw,dip}} \;\le\; (1+\eta)\, C_G^{\mathrm{raw,hap}}.$$

Therefore using $C_G^{\mathrm{raw,hap}}$ as the genomic budget throughout is conservative for both haploid and diploid organisms: a stricter functional estimate would move the threshold only toward coarser resolution, strengthening the impossibility conclusions, while a strict accounting of the second homolog could relax the bound by at most 1%. Even the narrowest threshold crossing reported below exceeds the plausible ploidy correction by at least an order of magnitude, so realistic ploidy adjustments cannot alter the qualitative conclusions. We also use "information quantity (bits)" as a neutral term in tables that mix Shannon and Hartley entropies, with the descriptor type specified per row: NSB Shannon estimate $H$ for empirical stochastic distributions, and zero-error addressability bound $H_0$ for combinatorial support counts.

**Corollary (Ensemble specification is forced).** *Below the threshold, biological function must be realized as a property of the conditional distribution $P(X_{\mathcal{C}'} \mid M, b)$ rather than as a property of individual realizations specified by the organism.*

*Proof.* By Theorem 1 part 2(b), for any strict sub-threshold refinement $\mathcal{C}'$ of some $\mathcal{C}_1^* \in \mathfrak{C}_1^*(B)$ with $H(X_{\mathcal{C}'} \mid b) > B$,

$$H(X_{\mathcal{C}'} \mid M, b) \;\ge\; H(X_{\mathcal{C}'} \mid b) - B \;>\; 0.$$

Hence $X_{\mathcal{C}'}$ is a nondegenerate random variable conditional on the organism's full specification $(M, b) = (g, U_{t_0:t_n}, b)$; its realized value is therefore not a deterministic function of that specification. Any biological function $F$ that the organism reliably produces at sub-threshold resolution must therefore take the same (or functionally equivalent) value across the ensemble of admissible realizations induced by $(M, b)$. Equivalently, $F$ is a functional of the conditional distribution $P(X_{\mathcal{C}'} \mid M, b)$ rather than a deterministic image of any single realization of $X_{\mathcal{C}'}$. □

Biology therefore necessarily operates as an ensemble-coded system at sub-threshold resolutions, with phenotypic precision arising from statistical regularities of the ensemble rather than from organism-programmed determination of individual molecular trajectories.

### A Information-theoretic notation and definitions

All random variables in this paper take values in finite sets. We collect the definitions of the information-theoretic quantities used throughout; standard references are Shannon (1948) and Cover & Thomas (2006).

**Shannon entropy.** For a random variable $X$ taking values in a finite set $\mathcal{X}$ with probability mass function $p(x) = \mathbb{P}(X = x)$,

$$H(X) \;=\; -\sum_{x\in\mathcal{X}} p(x)\log_2 p(x),$$

with the convention $0\log_2 0 = 0$. This is nonneg and satisfies $H(X) \le \log_2|\mathcal{X}|$, with equality iff $X$ is uniformly distributed.

**Conditional entropy.** For jointly distributed random variables $(X, Y)$ on $\mathcal{X} \times \mathcal{Y}$,

$$H(X \mid Y) \;=\; \sum_{y\in\mathcal{Y}} p(y)\, H(X \mid Y = y) \;=\; -\sum_{x,y} p(x, y)\log_2 p(x \mid y).$$

This is the expected remaining uncertainty in $X$ after observing $Y$. It satisfies $0 \le H(X \mid Y) \le H(X)$, with $H(X \mid Y) = 0$ iff $X$ is a.s. a deterministic function of $Y$.

**Mutual information.** The mutual information between $X$ and $Y$ is

$$I(X;Y) \;=\; H(X) - H(X \mid Y) \;=\; H(Y) - H(Y \mid X) \;=\; \sum_{x,y} p(x, y)\log_2 \frac{p(x, y)}{p(x)\,p(y)}.$$

It is symmetric, nonneg, and zero iff $X \perp\!\!\!\perp Y$. It satisfies $I(X;Y) \le \min\{H(X), H(Y)\}$. Conditional mutual information $I(X;Y \mid Z)$ is defined analogously by conditioning all terms on $Z$.

**Kullback–Leibler divergence.** For probability measures $\mu, \nu$ on a finite set $\mathcal{X}$,

$$D(\mu\|\nu) \;=\; \sum_{x\in\mathcal{X}} \mu(x)\log_2 \frac{\mu(x)}{\nu(x)},$$

with $D(\mu\|\nu) = +\infty$ if $\mu$ is not absolutely continuous w.r.t. $\nu$. It is nonneg (Gibbs' inequality) and zero iff $\mu = \nu$. It is not symmetric and does not satisfy the triangle inequality. Mutual information is a KL divergence: $I(X;Y) = D(P_{X,Y}\|P_X \otimes P_Y)$.

**Hartley entropy (zero-error support cost).** For a random variable $X$ on $\mathcal{X}$,

$$H_0(X) \;=\; \log_2|\operatorname{supp}(X)|, \qquad \operatorname{supp}(X) = \{x \in \mathcal{X} : p(x) > 0\}.$$

This is the number of bits required to address an arbitrary element of the support with zero error, regardless of the distribution. It satisfies $H(X) \le H_0(X)$, with equality iff $X$ is uniform on its support. The conditional Hartley entropy given a fixed value $z$ is $H_0(X \mid z) = \log_2|\operatorname{supp}(X \mid z)|$, where $\operatorname{supp}(X \mid z) = \{x : \mathbb{P}(X = x \mid Z = z) > 0\}$.

**Data-processing inequality.** If $X \to Y \to Z$ is a Markov chain (i.e., $X \perp\!\!\!\perp Z \mid Y$), then $I(X;Z) \le I(X;Y)$. In particular, if $Z = f(Y)$ for a deterministic function $f$, then $H(Z) \le H(Y)$ and $H(Z \mid X) \le H(Y \mid X)$.

##### Application to the biological framework

The random variables of the paper, all defined on the probability space of Appendix B, are:

- $g \in \mathcal{G}$: the genome (may be a fixed constant $g_0$ or a random variable over genomes);
- $U_{0:K-1} = (U_{\tau_0}, \dots, U_{\tau_{K-1}}) \in \mathcal{U}^K$: the discretized environmental signal;
- $W_{0:K-1} = (W_{\tau_0}, \dots, W_{\tau_{K-1}}) \in \mathcal{W}^K$: the thermal noise sequence (i.i.d.);
- $\mathbf{S} = (S_{\tau_0}, \dots, S_{\tau_K}) \in \Sigma^{K+1}$: the discretized trajectory, a deterministic function of $(g, U, W, S_{\tau_0})$;
- $X_\mathcal{C} = (\mathcal{C}(S_{\tau_0}), \dots, \mathcal{C}(S_{\tau_K})) \in \mathcal{C}^{K+1}$: the coarse-grained trajectory, a deterministic function of $\mathbf{S}$;
- $M = (g, U_{0:K-1})$: the organism-controlled "message" (the specification budget).

The key entropy quantities appearing in the main text are then:

- $H(X_\mathcal{C})$: the Shannon entropy of the coarse-grained trajectory. Finite since $X_\mathcal{C}$ takes values in the finite set $\mathcal{C}^{K+1}$.
- $H(X_\mathcal{C} \mid g, U)$: the conditional entropy of $X_\mathcal{C}$ given the genome and environment. This is the residual uncertainty in the trajectory that is *not* determined by the organism's specification. It equals zero iff the specification determines the trajectory a.s.
- $H_0(X_\mathcal{C} \mid z)$: the Hartley entropy of $X_\mathcal{C}$ conditioned on the fixed background $z = (g, U, \Phi)$. This is $\log_2$ of the number of distinct trajectories consistent with the specification — the "size of the fiber."
- $I(X_\mathcal{C}; M)$: the mutual information between the trajectory and the specification. Bounded above by $H(M) \le \log_2 |\mathrm{supp}(M)| \le C_G + C_E = B$.
- $D(\mu K \| \nu K)$: the KL divergence between two distributions after applying a Markov kernel $K$, used in the contraction hypothesis for Lemma 1b.

#### B Probability space and time discretization

We define the underlying probability space for the formal framework. Fix a genome length $n$, and let $\mathcal{G} = \{A, C, G, T\}^n$ be the set of possible genomes. Fix a temporal window $[t_0, t_n]$ and a discretization $t_0 = \tau_0 < \tau_1 < \cdots < \tau_K = t_n$ with step $\delta = \tau_{i+1} - \tau_i$.

The physical dynamics are modeled as a discrete-time Markov chain on $\Sigma$: at each step $i$, the transition

$$S_{\tau_{i+1}} \sim P(\cdot \mid S_{\tau_i}, g, U_{\tau_i})$$

is determined by the current state, the genome (which selects the transition kernel), and the current environmental input. Conditional on $(S_{\tau_i}, g, U_{\tau_i})$, the state $S_{\tau_{i+1}}$ is independent of all earlier states and of future inputs. The randomness in each transition is the thermal noise $W_{\tau_i}$, which is independent across time steps.

The underlying probability space is

$$(\Omega, \mathcal{F}, \mathbb{P}) = (\Sigma \times \mathcal{G} \times \mathcal{U}^K \times \mathcal{W}^K,\ 2^\Omega,\ \mathbb{P}),$$

where the first factor carries the initial state $S_{\tau_0}$, $\mathcal{U}$ is the (finite) set of discretized environmental inputs, $\mathcal{W}$ is the (finite) set of discretized noise values, and $\mathbb{P}$ is the joint distribution factoring as

$$\mathbb{P}(S_{\tau_0}, g, U_{0:K-1}, W_{0:K-1}) = P_0(S_{\tau_0} \mid g) \cdot P_{\mathcal{G}}(g) \cdot P_{\mathcal{U}}(U_{0:K-1}) \cdot \prod_{i=0}^{K-1} P_{\mathcal{W}}(W_i).$$

Here $P_0(\cdot \mid g)$ is the initial-state distribution (which may depend on the genome), $P_{\mathcal{G}}$ is a distribution over genomes (which may be a point mass at a specific genome $g_0$ for analyzing a single organism), $P_{\mathcal{U}}$ is the distribution of environmental signals, and $P_{\mathcal{W}}$ is the distribution of thermal noise (i.i.d. across steps). The trajectory $\mathbf{S} = (S_{\tau_0}, \ldots, S_{\tau_K})$ is then a deterministic function of $\omega = (S_{\tau_0}, g, U_{0:K-1}, W_{0:K-1})$: given the initial state and all inputs, each subsequent state is determined by the transition function $S_{\tau_{i+1}} = \Psi(S_{\tau_i}, g, U_{\tau_i}, W_{\tau_i})$ (we use $\Psi$ for the transition function to avoid confusion with $\Phi$, the universal compiler and physics substrate defined in the main text).

All Shannon entropies in the main text are computed with respect to this probability space. When the genome is held fixed at a specific value $g_0$ (as in the analysis of a single organism), all entropies are conditional on $g = g_0$, and the Shannon entropy $H(g)$ does not appear; only $C_E$ enters the Shannon bounds. The Hartley bounds use $C_G = \log_2 |\mathcal{G}|$ regardless of the distribution.

This discrete-time model is an approximation to the underlying continuous-time Markov chain (CTMC) dynamics of the physical system. A CTMC trajectory on $[t_0, t_n]$ is determined by the initial state, the (random) number of jumps $N$, the jump times $\tau_1, \ldots, \tau_N \in [t_0, t_n]$, and the sequence of visited states. Because the jump times are continuous random variables, the full CTMC trajectory has infinite Shannon entropy, and the discrete approximation is essential for the entropy comparisons in the main text to be well-defined. The impossibility results hold for every discretization step $\delta > 0$: refining the discretization can only increase the entropy $H(X_{\mathcal{C}})$ (since finer temporal resolution reveals more of the trajectory), so any impossibility at resolution $\delta$ implies impossibility at all finer resolutions.

### C Connection between Lemma 1 and CTMC dynamics

Lemma 1 is stated for an abstract algorithm reading i.i.d. fair bits. We connect this to the discretized Markov chain as follows.

At each step $i$, the transition $S_{\tau_{i+1}} \mid (S_{\tau_i} = s, g, U_{\tau_i} = u)$ has a conditional distribution $P(\cdot \mid s, g, u)$ on $\Sigma$. This distribution can be sampled using the inverse-CDF method: draw a uniform random variable $V_i \in [0, 1)$ and apply the quantile function $F^{-1}_{s,g,u}(V_i)$. Since $|\Sigma|$ is finite, sampling $V_i$ to sufficient precision requires at most $\lceil \log_2 |\Sigma| \rceil$ fair bits (via a rejection-sampling or arithmetic-coding construction; see Cover & Thomas 2006, Chapter 5).

Over $K$ steps, the total fair bits consumed is $T \le K \lceil \log_2 |\Sigma| \rceil$, which is finite. The output is the discretized trajectory $\mathbf{S} = (S_{\tau_0}, \ldots, S_{\tau_K}) \in \Sigma^{K+1}$. Lemma 1 then applies with $X = \mathbf{S}$, giving

$$\mathbb{E}[T \mid g, U_{t_0:t_n}] \ge H(\mathbf{S} \mid g, U_{t_0:t_n}).$$

The thermal noise $W_{0:K-1}$ of the physical system corresponds to the fair bits consumed: it is the randomness source from which each transition draws the information not determined by $(g, U)$.

### D Hartley and Shannon capacity

We clarify the relationship between the two capacity notions used in the main text.

**Hartley capacity.** For a message $M$ taking values in a finite set $\mathcal{M}$, the Hartley entropy is $H_0(M) = \log_2 |\mathcal{M}|$. This is the number of bits required to address an arbitrary element of $\mathcal{M}$ with zero error. It does not depend on any probability distribution.

**Shannon capacity.** The Shannon entropy $H(M) = -\sum_{m\in\mathcal{M}} P(m) \log_2 P(m)$ depends on the distribution $P$ and satisfies $H(M) \le H_0(M) = \log_2 |\mathcal{M}|$, with equality if and only if $P$ is uniform on $\mathcal{M}$.

**Relationship to the specification budget.** For the genome, $\mathcal{G} = \{A, C, G, T\}^n$, so $C_G = H_0(g) = 2n$ bits. For a specific organism with genome $g_0$, the genome is a constant (a point mass), so $H(g) = 0$. The quantity $C_G = 2n$ is therefore the Hartley capacity, not the Shannon entropy.

In the main theorem:

- **Prong (2a)** uses only $C_G$ as a Hartley capacity: a string of $2n$ bits can address at most $2^{2n} = 4^n$ distinct objects. This is a combinatorial fact requiring no probability model.
- **Prong (2b)** uses the bound $H(M) \le \log_2 |\operatorname{supp}(M)| \le B = C_G + C_E$. This holds for *any* distribution on $M$, including a point mass on a specific genome (in which case $H(M) = H(U) \le C_E$ and the bound with $B = C_G + C_E$ is looser but still valid). The Shannon prong is therefore a quantitative refinement of the Hartley prong: it gives a tighter lower bound on the runtime randomness consumed, at the cost of requiring a probability model.

For a single organism with a fixed genome $g_0$, the Shannon prong simplifies: $H(X_{\mathcal{C}'} \mid g_0, U, b) \ge H(X_{\mathcal{C}'} \mid g_0, b) - C_E$. The genome enters the Shannon picture by selecting the transition kernel (different $g$ gives different Markov chains), not as a random information source. The Hartley prong, by contrast, works over the space of all possible genomes and requires no such distinction.

### E Contraction assumption for driven chains

The uniform KL contraction hypothesis used in Lemma 1b is

$$D(\mu K_{i,j}^{g,U} \| \nu K_{i,j}^{g,U}) \le e^{-(j-i)\delta/\tau_{\text{mix}}} D(\mu \| \nu)$$

for all probability measures $\mu, \nu$ on $\Sigma$ and all steps $j \ge i$, where $K_{i,j}^{g,U}$ is the multi-step Markov kernel from step $i$ to step $j$ conditional on $(g, U_{i:j-1})$, and $\delta$ is the time-discretization step.

For a **time-homogeneous** finite irreducible aperiodic chain, uniform KL contraction holds with a rate determined by the modified log-Sobolev constant (MLSI) of the single-step kernel. This is implied by, but weaker than, the spectral gap. The “uniform” qualifier (contraction for all pairs $\mu, \nu$, not only for $\mu$ vs. the stationary distribution $\pi$) is a Dobrushin-type condition equivalent to uniform ergodicity on finite state spaces.

For a **driven** (time-inhomogeneous) chain, the single-step kernels $P(\cdot \mid \cdot, g, U_{\tau_i})$ depend on the time-varying environmental input. The contraction hypothesis requires that each single-step kernel satisfies KL contraction and that the contraction rate can be chosen uniformly over the range of environmental inputs encountered. In terms of the underlying continuous-time dynamics with instantaneous generators $Q(g, U_{\tau_i})$, sufficient conditions include:

1. Each $Q(g, U_{\tau_i})$ has a strictly positive diagonal (no absorbing states), ensuring irreducibility at each step.

2. The minimum transition rate is bounded below uniformly over all environmental inputs, giving a uniform lower bound on the per-step contraction rate.

3. The environmental signal varies slowly relative to $\tau_{\text{mix}}$, so that the chain approximately equilibrates between environmental changes.

These conditions are physically reasonable for biological systems at molecular scales in thermal equilibrium or near it: the thermal bath ensures that every molecular configuration is reachable from every other (condition 1), the thermal collision rate provides a lower bound on transition rates (condition 2), and molecular relaxation (∼ns) is fast relative to environmental variation (∼ms–s) (condition 3). The slow/fast decomposition in the main text restricts the contraction hypothesis to the fast residual variables $Z_{\tau_i}$, for which these conditions are most easily verified.

### F Precise statement of Lemma 2

The main-text statement of Lemma 2 uses measurable functions to define "current function" and "forward program." We record here the precise conditional-independence structure that underlies it.

**Setup.** Let $(S_k)_{k=0}^{n}$ be a discrete-time Markov chain on $\Sigma$, driven by environmental inputs $(U_k)_{k=0}^{n-1}$ and noise $(W_k)_{k=0}^{n-1}$, with transition

$$S_{k+1} = \Psi(S_k, U_k, W_k)$$

for some deterministic function $\Psi : \Sigma \times \mathcal{U} \times \mathcal{W} \to \Sigma$. The Markov property gives the conditional independence

$$(S_0, \ldots, S_{k-2}, g) \perp\!\!\!\perp (S_k, S_{k+1}, \ldots, S_n) \mid (S_{k-1}, U_{k-1:n-1}, W_{k-1:n-1}).$$

**Definition.** A *programmed forward action from step k* is any measurable function $A = \phi(S_{k-1}, U_{k:n})$ that depends on the current state and future environmental inputs but not on future noise $W_{k:n}$. This captures the idea that a "programmed" outcome is one determined by the organism's specification stores $(S_{k-1}, g, U)$, where $g$ is encoded in $S_{k-1}$ via its influence on the transition kernel. Future noise $W_{k:n}$ is by definition not programmed.

**Lemma 2 (precise form).** For any measurable $\phi : \Sigma \times \mathcal{U}^{n-k+1} \to \mathcal{A}$,

$$H(\phi(S_{k-1}, U_{k:n}) \mid U_{k:n}) \leq H(S_{k-1} \mid U_{k:n}) \leq \log_2 |\Sigma|.$$

*Proof.* The first inequality is the data-processing inequality: $\phi(S_{k-1}, U_{k:n})$ is a deterministic function of $(S_{k-1}, U_{k:n})$, so $H(\phi(S_{k-1}, U_{k:n}) \mid U_{k:n}) \leq H(S_{k-1} \mid U_{k:n})$. The second inequality is $H(S_{k-1} \mid U_{k:n}) \leq H(S_{k-1}) \leq \log_2 |\Sigma|$. □

The lemma says that current function and forward program must share the same $\log_2 |\Sigma|$-bit budget. Any information devoted to future instructions must be physically present in the current state, reducing the capacity available for current physiological function. This is a finite-state bottleneck, not a dynamical claim: it holds regardless of the specific transition kernel, the mixing rate, or the nature of the environmental signal.

### Supplementary Note 2: Environmental Rescue Bounds

#### S1. What counts as an environmental rescue channel

The environmental channel in the main text is a *programmatic* channel: an exogenous process that carries decodable information into the biological system. It is not synonymous with the background physical conditions under which biology occurs. The universal compiler and physics substrate — the laws of physics, solvent, thermal bath, hydrophobic effect, ordinary chemistry, and other invariant background conditions — are required for biological organization, but they are not message channels carrying organism-specific bits.

Let $\mathfrak{E}_{\mathrm{bio}}$ denote the class of biologically admissible environmental channels over a window $[t_0, t_n]$. A channel $\mathcal{E} \in \mathfrak{E}_{\mathrm{bio}}$ consists of an external message $M_E$, a causal encoder producing a boundary process $E_{t_0:t_n}$, receiver variables $R_{t_0:t_n} \subseteq S_{t_0:t_n}$, and a decoder producing $\widehat{M}_E$. Its transmitted specifying information is $I(M_E; \widehat{M}_E)$. The integrated programmatic environmental capacity is

$$C_E([t_0, t_n]) = \sup_{\mathcal{E} \in \mathfrak{E}_{\mathrm{bio}}} \sup_{P(M_E),\, \mathrm{enc},\, \mathrm{dec}} I_{\mathcal{E}}(M_E; \widehat{M}_E), \tag{S1}$$

where the supremum is restricted by the physical limits of the modality: rate, fidelity, addressability, synchronization, and biocompatible actuation. If a channel has an instantaneous addressed information-rate envelope $r_{\mathcal{E}}(t)$, then

$$C_E([t_0, t_n]) \le \sup_{\mathcal{E} \in \mathfrak{E}_{\mathrm{bio}}} \int_{t_0}^{t_n} r_{\mathcal{E}}(t)\, dt. \tag{S2}$$

The adjective *addressed* is important. A broadcast field contributes one shared low-dimensional input unless it can deliver distinct decodable messages to distinct target cells or subcellular regions. A signal that changes ensemble dynamics but carries no decodable microstate-specific message is a boundary condition, not a deterministic microstate program. For the natural-rescue claim below, $\mathfrak{E}_{\mathrm{bio}}$ denotes naturally realized biological source–receiver architectures. Engineered external controllers are not ruled out; they enlarge the specification system, and their sensing, memory, communication, and actuation capacity must be included in the enlarged $C_E$.

#### S2. Required rescue rate

Let

$$B_0 = C_G + C_E^{\text{already counted}}$$

denote the organism-controlled budget already included in the model, and let $b$ denote the fixed non-message background. For deterministic microstate rescue, the remaining zero-error addressability deficit is

$$\Delta_0 = \left[H_0(X_{\mathcal{C}'} \mid b) - B_0\right]_+, \tag{S3}$$

and for stochastic sampling rescue the corresponding Shannon deficit is

$$\Delta_1 = \left[H(X_{\mathcal{C}'} \mid b) - B_0\right]_+, \tag{S4}$$

where $[x]_+ = \max\{x, 0\}$. Equivalently, after conditioning on the already-counted message, the residuals are $H_0(X_{\mathcal{C}'} \mid g, U, b)$ and $H(X_{\mathcal{C}'} \mid g, U, b)$, with no further subtraction of $B_0$. A new channel acting over a time window $T$ must supply at least

$$R_{\text{req}} = \Delta/T \tag{S5}$$

addressed bits per second, where $\Delta$ is $\Delta_0$ or $\Delta_1$ depending on whether the claim is zero-error specification or runtime sampling.

For the representative bacterial case used in the main text, the conservative unresolved gap is $\Delta_0 \sim 5 \times 10^6$ bits, $T_{\text{gen}} = 1800$ s, so

$$R_{\text{req}}^{\text{bac}} \approx 2.78 \times 10^3 \ \text{bits}\,\text{s}^{-1}\,\text{cell}^{-1}. \tag{S6}$$

For a deliberately conservative eukaryotic comparison retaining the same $5 \times 10^6$-bit gap over a 24-hour cell cycle, $T_{\text{gen}} = 8.64 \times 10^4$ s,

$$R_{\text{req}}^{\text{euk}} \approx 5.8 \times 10^1 \ \text{bits}\,\text{s}^{-1}\,\text{cell}^{-1}. \tag{S7}$$

This eukaryotic number is not intended as a realistic full-cell gap; it is a permissive lower benchmark. Larger eukaryotic state spaces only strengthen the conclusion.

#### S3. Feedback-or-override dichotomy

Let $\Omega_t$ be the set of current microstates still admissible immediately before an environmental symbol $E_t = e_t$ is applied:

$$\Omega_t = \text{supp}(S_t \mid g, U_{t_0:t}, E_{t_0:t^-}). \tag{S8}$$

**Proposition 3** (Feedback-or-override dichotomy)**.** *Assume* $|\Omega_t| > 1$*, and suppose the rescue objective is a prescribed common next state* $s^\star$ *(or prescribed target equivalence class). If the rescue symbol* $E_t$ *is open-loop,*

$$E_t \perp S_t \mid g, U_{t_0:t}, E_{t_0:t^-},$$

*then zero-error rescue at the next step is possible only if the actuation is state-erasing on* $\Omega_t$*:*

$$K_t(\cdot \mid s, e_t, g, U_t) = \delta_{s^\star} \quad \textit{for all } s \in \Omega_t. \tag{S9}$$

*If the actuation is not state-erasing, deterministic rescue requires a feedback observation* $O_t$ *that distinguishes the control-relevant equivalence class* $C_t = C(S_t)$*. Consequently,*

$$I(O_t; S_t \mid g, U_{t_0:t}, E_{t_0:t^-}) \geq H(C_t \mid g, U_{t_0:t}, E_{t_0:t^-}), \tag{S10}$$

*and in the zero-error equiprobable case it requires at least* $\log_2 |\Pi_t|$ *bits, where* $\Pi_t$ *is the partition of* $\Omega_t$ *into control-relevant classes.*

*Proof.* If the same open-loop symbol maps two admissible states to different next-state distributions, then the next state has residual uncertainty after conditioning on $(g, U, E)$. Zero residual uncertainty for a prescribed common target therefore requires all states in $\Omega_t$ to be collapsed to that target, i.e. reset/override. If override is not available, the correct actuation depends on which control-relevant class the current state occupies. Any observation supporting a deterministic control law must recover that class. Since $C_t$ is a deterministic function of $S_t$, data processing gives the stated mutual-information requirement. □

Thus a deterministic environmental rescue must either measure the present microstate and send back a tailored signal, or erase the current state. Ordinary biological cues do neither at microstate-resolved precision; they bias the ensemble.

#### S4. Chemical diffusive signaling

Consider one ligand species sensed by a receptor of effective radius $a$, concentration $c$, and diffusion coefficient $D$. Berg–Purcell counting (Berg & Purcell, 1977) gives

$$\mathrm{SNR}^2 \leq 4\pi DcaT \tag{S11}$$

over integration time $T$. An optimistic scalar Gaussian-equivalent conversion gives the proxy

$$I_T^{(\mathrm{G})} = \tfrac{1}{2}\log_2(1 + 4\pi DcaT), \tag{S12}$$

and hence

$$R_{\mathrm{chem}}^{(\mathrm{G})}(T) = \frac{1}{2T}\log_2(1 + 4\pi DcaT). \tag{S13}$$

For a diffusive symbol originating a distance of order $\ell_{\mathrm{cell}}$ away, transport requires

$$\sqrt{DT} \gtrsim \ell_{\mathrm{cell}} \quad \Longrightarrow \quad T \gtrsim \ell_{\mathrm{cell}}^2/D. \tag{S14}$$

Using this fastest cell-scale diffusive refresh gives

$$R_{\mathrm{chem}}^{(1)} \approx \frac{D}{2\ell_{\mathrm{cell}}^2}\log_2(1 + 4\pi ca\ell_{\mathrm{cell}}^2). \tag{S15}$$

For bacterial-scale values $D = 100\ \mu\mathrm{m}^2\mathrm{s}^{-1}$, $c = 100$ nM $\simeq 60\ \mu\mathrm{m}^{-3}$, $a = 1$ nm $= 10^{-3}\ \mu$m, and $\ell_{\mathrm{cell}} = 1\ \mu$m, we obtain

$$R_{\mathrm{chem}}^{(1)} \approx 40.6\ \mathrm{bits\,s}^{-1}. \tag{S16}$$

At $c = 1\ \mu$M, this becomes

$$R_{\mathrm{chem}}^{(1)} \approx 155\ \mathrm{bits\,s}^{-1}. \tag{S17}$$

Thus, under the stated 1-nm effective-capture-radius model, the bacterial rescue benchmark requires roughly $2.78 \times 10^3/40.6 \approx 68$ independent ligand channels at 100 nM, or $\approx 18$ at 1 $\mu$M, assuming orthogonality, independence, fast receptor reset, and perfect downstream decoding.

For a eukaryotic-scale cell, $\ell_{\mathrm{cell}} = 10\ \mu$m, the same proxy yields

$$R_{\mathrm{chem}}^{(1)} \approx 3.13\ \mathrm{bits\,s}^{-1}\quad (100\ \mathrm{nM}), \qquad R_{\mathrm{chem}}^{(1)} \approx 4.78\ \mathrm{bits\,s}^{-1}\quad (1\ \mu\mathrm{M}). \tag{S18}$$

These parameter-conditioned estimates still ignore the central feedback problem: ligand concentration reports a low-dimensional external field, not the receiver's current microstate. A deterministic chemical rescue would require a measurement channel from the cell interior plus a return chemical channel, or a destructive state-erasing chemical override. Natural chemical signaling is therefore an ensemble-biasing mechanism, not a microstate instruction tape.

#### S5. Mechanical and acoustic signaling

For far-field wave-based addressing of a cell of size $\ell_{\mathrm{cell}}$ with a classical acoustic wave of speed $c_s$, the carrier wavelength must satisfy $\lambda \lesssim 2\ell_{\mathrm{cell}}$, hence

$$f_{\mathrm{addr}} \gtrsim \frac{c_s}{2\ell_{\mathrm{cell}}}. \tag{S19}$$

With $c_s \simeq 1.5 \times 10^3 \ \mathrm{m\,s^{-1}}$,

$$f_{\mathrm{addr}}^{\mathrm{bac}} \gtrsim 7.5 \times 10^8 \ \mathrm{Hz}, \qquad f_{\mathrm{addr}}^{\mathrm{euk}} \gtrsim 7.5 \times 10^7 \ \mathrm{Hz}. \tag{S20}$$

At such frequencies, acoustic attenuation in water or tissue is substantial, and biological receivers are low-dimensional mechanosensors (tension, strain, pressure), not high-dimensional intracellular actuators.

A finite acoustic capacity requires an amplitude bound. If $p_{\max}$ is a biocompatible pressure amplitude, the acoustic intensity is

$$I_{\max} = \frac{p_{\max}^2}{2\rho c_s}, \tag{S21}$$

so the received power on cross-section $A \sim \ell_{\mathrm{cell}}^2$ is

$$P_r \lesssim \frac{p_{\max}^2}{2\rho c_s} \ell_{\mathrm{cell}}^2 G_{\mathrm{ac}}(f, L), \tag{S22}$$

where $G_{\mathrm{ac}}(f, L) \leq 1$ is the frequency- and path-dependent propagation gain. In the low-SNR thermal limit,

$$C_{\mathrm{ac}} \lesssim \frac{P_r}{k_B T \ln 2}. \tag{S23}$$

For $p_{\max} = 1$ Pa, $\rho = 10^3 \ \mathrm{kg\,m^{-3}}$, $T = 310$ K, and representative path lengths $L_{\mathrm{bac}} = 100 \ \mu$m and $L_{\mathrm{euk}} = 1$ cm, representative attenuation laws give single-cell rates of order 1–10 bits $\mathrm{s^{-1}}$. These are parameter-conditioned examples, not amplitude-independent modality bounds. Without an amplitude or power bound, no finite amplitude-independent Shannon upper bound exists for acoustic waves. The unconditional obstruction is addressability plus the lack of a high-dimensional state-reading receiver.

#### S6. Electrical signaling

In physiological saline, static electrostatic fields are screened over the Debye length, typically below a nanometre. The generic electrical degree of freedom that survives at the cellular boundary is the membrane voltage or junctional current, a low-dimensional collective variable.

A membrane or junction can be modeled as a single scalar RC channel with

$$\tau_m = R_m C_m, \qquad B_{\mathrm{elec}} \lesssim \frac{1}{2\pi \tau_m}. \tag{S24}$$

Johnson–Nyquist noise (Johnson, 1928; Nyquist, 1928) over bandwidth $B$ has

$$V_n^2 = 4 k_B T R_m B. \tag{S25}$$

Thus, for voltage swing $V_{\max}$,

$$C_{\mathrm{elec}} \leq B_{\mathrm{elec}} \log_2\left(1 + \frac{V_{\max}^2}{4 k_B T R_m B_{\mathrm{elec}}}\right). \tag{S26}$$

With $R_m = 10^9 \ \Omega$, $V_{\max} = 100$ mV, $\tau_m = 10$ ms for a bacterial-scale interface and 100 ms for a eukaryotic unspecialized membrane, one obtains rates of order $4 \times 10^2$ bits $\mathrm{s^{-1}}$ and $4.5 \times 10^1$ bits $\mathrm{s^{-1}}$, respectively. These are generic single-interface estimates. Specialized natural interfaces can be larger, and dense electrode arrays or patch-clamp feedback systems are external controllers rather than ordinary environmental rescue channels.

#### S7. Electromagnetic and optical signaling

Far-field diffraction-limited addressing requires

$$\lambda \lesssim 2\ell_{\text{cell}} \quad \Longleftrightarrow \quad f \gtrsim \frac{c}{2\ell_{\text{cell}}}. \tag{S27}$$

Thus

$$f_{\text{addr}}^{\text{bac}} \gtrsim 1.5 \times 10^{14}\ \text{Hz}, \qquad f_{\text{addr}}^{\text{euk}} \gtrsim 1.5 \times 10^{13}\ \text{Hz}. \tag{S28}$$

Propagation through tissue obeys Beer–Lambert attenuation,

$$I(L) = I_0 e^{-\mu_{\text{eff}} L}. \tag{S29}$$

Even in favorable optical windows, centimeter-scale depths produce strong attenuation; at water absorption bands, the penalty is larger. A classical optical channel has no finite amplitude-independent Shannon capacity: capacity grows with received power. Therefore the rigorous general statements are far-field addressability, attenuation, and the absence of a natural intracellular high-dimensional optical decoder. Engineered optogenetic or microscope-controlled systems can deliver control by adding external source memory, sensors, computation, and actuation; they enlarge the specification system rather than evade the bound.

#### S8. Thermal signaling

Temperature fields obey diffusion with thermal diffusivity $\alpha_{\text{th}}$. A cell-scale relaxation bandwidth obeys

$$B_{\text{th}} \lesssim \frac{\alpha_{\text{th}}}{\ell_{\text{cell}}^2}. \tag{S30}$$

For water-like $\alpha_{\text{th}} \approx 1.4 \times 10^{-7}\ \text{m}^2\,\text{s}^{-1}$,

$$B_{\text{th}}^{\text{bac}} \lesssim 1.4 \times 10^5\ \text{s}^{-1}, \qquad B_{\text{th}}^{\text{euk}} \lesssim 1.4 \times 10^3\ \text{s}^{-1}. \tag{S31}$$

Thermal fluctuations in volume $V \sim \ell^3$ satisfy

$$\sigma_T^2 = \frac{k_B T^2}{\rho c_p V}. \tag{S32}$$

Rate alone is not the primary obstruction: forced local heating can transmit energy rapidly. The obstruction is that temperature is a broadcast scalar field with no generic cell-specific addressing or high-dimensional state-sensitive receiver. Thermal cues can bias ensembles and alter rates; they cannot encode a deterministic molecular instruction tape without external microheaters and feedback instrumentation.

#### S9. Quantum signaling

Holevo's bound (Holevo, 1973) limits $n$ transmitted qubits to at most $n$ classical bits of accessible information in the absence of prior entanglement. Closing a gap $\Delta_0$ therefore requires at least

$$n_Q \geq \Delta_0 \tag{S33}$$

quantum carriers at the information-accounting level. Quantum encoding therefore does not remove the deficit. A biological quantum rescue would additionally require a source, protected channel, receiver, readout, and molecular actuator at the target resolution. Decoherence in warm biological media constrains coherent storage and processing, but does not by itself imply a universal transmission-rate bound. If the quantum state is immediately measured and used as a classical signal, the problem reduces to the classical communication-and-control channels above and gains no information-theoretic escape.

#### S10. Source and receiver requirements

A successful deterministic rescue source–receiver pair would have to satisfy all of the following:

1. **Rate:** deliver $\Delta/T$ addressed bits per target during the control window.
2. **Addressability:** deliver distinct symbols to the relevant cells or subcellular regions; a shared broadcast field is not enough.
3. **Fidelity:** maintain sufficiently low cumulative error over the full trajectory.
4. **Synchronization:** deliver symbols before the targeted degrees of freedom mix or diverge.
5. **State access or state erasure:** either measure the current microstate at control-relevant resolution or erase it by override.
6. **Actuation depth:** couple the received symbols to the specific molecular degrees of freedom, not just to a low-dimensional receptor, voltage, or stress summary.
7. **Source memory:** store the desired trajectory and, in feedback mode, the measured current state in an external controller.

No known naturally realized biological source–receiver architecture satisfies all of these requirements at microstate resolution. Purely technological external controllers are not ruled out; if they supply the missing sensing, memory, communication, and actuation, they enlarge the controlled system and their information is counted in the enlarged $C_E$.

#### S11. Cell-to-cell signaling and isogenic no-bootstrap

A natural rescue proposal is that other cells supply the missing information. Non-override rescue must cooperate with the receiver's genome-dependent response map. Thus the source must know a genome-response sufficient statistic of the receiver, or be isogenic with it. But isogenic copies do not multiply genomic information. When $g$ ranges over the admissible genome alphabet and $C_G = H_0(g)$ denotes its Hartley capacity, if $g^{(1)} = \cdots = g^{(m)} = g$ almost surely, then

$$H_0(g^{(1)}, \ldots, g^{(m)}) = H_0(g) = C_G, \qquad H(g^{(1)}, \ldots, g^{(m)}) = H(g) \leq C_G. \tag{S34}$$

A population of isogenic cells can amplify, route, and redundantly encode the same genome-specific information, but it cannot create new genome-specific program bits by replication alone. Additional information in cell–cell signals must come from sender microstates, exogenous inputs, or runtime fluctuations; these are ensemble dynamics, not deterministic microstate-program bits stored by the shared genome.

#### S12. Repeated deterministic genome draws do not amplify information

Another loophole is that the cell might repeatedly query or recombine pieces of the genome to generate a long deterministic instruction stream. Let $Y_{t_0:t_n}$ be the complete transcript produced by any deterministic genome-query scheme, including adaptive index choices, retrieved substrings, and emitted symbols. Conditional on initial state and environmental input, the transcript is a deterministic function of $g$, hence

$$H(Y_{t_0:t_n} \mid S_{t_0}, U_{t_0:t_n}) \leq H(g \mid S_{t_0}, U_{t_0:t_n}) \leq C_G. \tag{S35}$$

Deterministic recombination of a static string cannot create more organism-specific information than the string contains. If fresh randomness or current-state information is injected, the excess is counted by the runtime-randomness lemma or by causal locality, not by the genome.

#### S13. Modality summary

Chemical diffusion supplies the cleanest quantitative natural-channel example and, under the stated single-receptor-scale assumptions, requires multiple independent addressed ligand channels for a single bacterial-scale rescue. Mechanical, electrical, optical, and thermal channels are constrained by addressability, attenuation, low-dimensional receiver structure, or the need for explicit power bounds. Quantum encoding does not remove the information deficit and lacks a known natural microstate-resolved source–receiver architecture. Across modalities, the same structural conclusion holds: natural environmental channels can bias and coordinate ensembles at functional coarse-grainings, but no known naturally realized continuous environmental channel acts as a deterministic, addressable microstate instruction tape (Shannon, 1956). Engineered external controllers are not ruled out; when present, they enlarge the specification system and must pay the missing information cost in $C_E$.

#### S14. Modality summary table

#### S15. What belongs to $C_E$ — objections and replies

The modality survey above bounds candidate natural channels one at a time. The conclusion does not depend on the survey being exhaustive: any proposed environmental rescue must carry decodable information about the target biological outcome and satisfy the source–receiver requirements of Sections S3 and S10. The following objections make this distinction explicit.

**First objection — the raw environment is information-rich.** *The environment contains astronomically many physical bits and can therefore supply whatever the genome does not specify.* Gross physical complexity is not organism-specific specification. The relevant quantity is decodable information about the target outcome, not the number of microscopic degrees of freedom in the surroundings. The same soil, light, water, and climate can support many different organisms, while a given lineage can develop across a broad permissive range of these conditions. Thus the ambient conditions alone do not identify which organism is present. A recovered diagnostic genomic sequence, by contrast, can identify a lineage. On a lifeless but habitable exoplanet, a sequence diagnostic of *Lepidochelys kempii* would support the retrodiction that this lineage had been present,

| Modality | Key relation | Bacterial example | Eukaryotic example | Status |
|---|---|---|---|---|
| Chemical diffusion | $R^{(1)}_{\mathrm{chem}} \approx \frac{D}{2\ell^2}\log_2(1+4\pi ca\ell^2)$ | 40.6 bps at 100 nM, $a = 1\,\mathrm{nm}$ | 3.13 bps at 100 nM, $a = 1\,\mathrm{nm}$ | Parameter-conditioned Gaussian-equivalent proxy; low-dimensional and lacks microstate feedback/actuation |
| Mechanical / acoustic | $C_{\mathrm{ac}} \lesssim P_r/(k_B T \ln 2)$ with $P_r$ from (S22) | $\sim$ 1–10 bps for $p_{\max} = 1\,\mathrm{Pa}$, $L = 100\,\mu\mathrm{m}$ | $\sim$ 1–10 bps for $p_{\max} = 1\,\mathrm{Pa}$, $L = 1\,\mathrm{cm}$ | Parameter-conditioned; no finite amplitude-independent bound, and no natural high-dimensional receiver |
| Electrical | $C_{\mathrm{elec}} \le B\log_2(1 + V^2_{\max}/(4k_B T R_m B))$ | $4.0 \times 10^2$ bps | $4.5 \times 10^1$ bps | Generic single-interface estimate; specialized or engineered interfaces can be larger |
| EM / optical | $f_{\mathrm{addr}} \ge c/(2\ell)$, $I(L) = I_0 e^{-\mu_{\mathrm{eff}} L}$ | $f_{\mathrm{addr}} \gtrsim 1.5 \times 10^{14}$ Hz | $f_{\mathrm{addr}} \gtrsim 1.5 \times 10^{13}$ Hz | Far-field addressing and depth limited; no finite amplitude-independent capacity bound without power/receiver assumptions |
| Thermal | $B_{\mathrm{th}} \lesssim \alpha_{\mathrm{th}}/\ell^2$ | $B_{\mathrm{th}} \lesssim 1.4 \times 10^5\ \mathrm{s}^{-1}$ | $B_{\mathrm{th}} \lesssim 1.4 \times 10^3\ \mathrm{s}^{-1}$ | Relaxation can be fast; natural temperature fields lack generic per-cell microstate addressability |
| Quantum | $n_Q \ge \Delta_0$ | $\ge 5 \times 10^6$ carriers for the benchmark gap | Same benchmark if $\Delta_0$ is retained | No information advantage; coherence adds implementation constraints, and no natural microstate-resolved architecture is known |

Table S1: **Environmental-rescue constraints across signaling modalities.** The numerical entries are parameter-conditioned examples rather than universal modality capacities. The general natural-world obstruction is the conjunction of rate, addressability, fidelity, synchronization, microstate access or state erasure, and actuation depth. Engineered external controllers are not ruled out; they enlarge the specification system and their information belongs in the enlarged $C_E$.

whereas beach temperatures compatible with its development would not: the same temperature regime is compatible with many lineages. Environmental bit-richness enters $C_E$ only to the extent that an admissible measurement and decoder extract organism-specific information from it.

**Second objection — temperature determines sex.** *Temperature-dependent sex determination in* L. kempii *shows that the environment specifies phenotype.* It shows that an environmental signal can select among coarse biological branches, which the framework already includes in $C_E$ (Geis *et al.*, 2005). Conditional on a genome that installs a temperature-sensitive developmental network and the alternative sex-specific generators, incubation temperature can carry at most one bit about a binary sex outcome,

$$I(\text{sex}; \text{temperature} \mid g) \leq 1 \text{ bit},$$

with equality approached only for a reliable binary selector. The same temperature does not identify the lineage and has no sex-determining meaning in the absence of the biological decoder. Such signals are therefore low-dimensional selectors among genome-conditioned developmental branches above $\mathcal{C}^*$, not microstate blueprints.

**Third objection — the channel survey is incomplete.** *The rescuing channel is simply one that the finite survey did not examine.* Any unlisted channel must still create conditional dependence between the target observable and an environmental input at fixed genotype. If

$$I(X_{\mathcal{C}}; U \mid g) = 0,$$

then no decoder can extract organism-specific specifying information from $U$ for that target. Environmental and $G \times E$ variance components are empirical diagnostics of such dependence, although they are not complete measures of mutual information. The modality calculations are therefore corroborative rather than logically exhaustive: a proposed omitted channel must identify a source, receiver, and target-dependent signal carrying the missing information, rather than appeal to unspecified environmental complexity.

**Fourth objection — another substrate is the hidden blueprint.** *A bioelectric prepattern, morphogenetic field, structured solvent, or other substrate carries the organism-specific plan.* The candidate falls into one of the categories already defined. If it varies with organismal identity or target outcome at fixed genome and is decodable by the system, it is a message channel and contributes to $C_E$; if it is physically instantiated in the inherited or current state, it is state-carried information counted by causal locality; if it is shared across organisms, it belongs to the fixed compiler/substrate $\Phi$; and if its fine realization is supplied by uncontrolled fluctuations, it belongs to $W$. A candidate cannot be both organism-specific and independent of every organism-specific source. The point is not to deny bioelectric, mechanical, or field-level causation, but to prevent it from being introduced as an uncounted information store.

#### S16. The oocyte as inherited state, not a second specification — objections and replies

The oocyte contains extensive organized material inherited from the maternal cell. The question is not whether this state matters — it plainly does — but whether its realized organization constitutes a second, independently encoded organism-specific blueprint rather than continued physical state.

**First objection — a second blueprint beside DNA.** *The oocyte carries an organism-specific program in addition to the genome and thereby specifies what DNA leaves undetermined.* If such a program is claimed to close the full residual fine-state deficit, it must carry at least the corresponding residual addressability. Naming that store "the oocyte" does not remove the information requirement; it relocates it and raises the same questions of origin, copying fidelity, and maintenance. Low-capacity maternal determinants and state variables are fully compatible with the framework, but a proposed genome-scale second blueprint must be exhibited as an independently decodable message and counted explicitly.

**Second objection — realized content is itself a compressed specification.** *The egg contains a large amount of organism-matched structure; therefore that structure must be a stored program for the offspring.* Exact agreement between parent and offspring state does not imply that the generative program contained the realized bits. Consider a fixed program on a universal computer that receives $N$, draws a string

$$R \leftarrow \text{RandomBits}(N),$$

from an external random-bit source, and then makes an exact copy

$$R' \leftarrow R.$$

The program has a description length independent of the realized $N$-bit string (apart from specifying $N$), yet $R' = R$ bit for bit. This is a general fact of computation, independent of programming language or hardware, and it remains true if the original program and machine are later destroyed or forgotten: the surviving matching strings are copied realized state, not evidence that the program contained the particular draw or could regenerate it de novo. Biologically, genome-conditioned physical dynamics and runtime fluctuations generate a maternal cell state, and cell division transmits much of that realized state by physical continuity. The presence of extensive matched state in the oocyte therefore does not by itself establish a second compressed blueprint; one must identify a separate message and decoder rather than the continued state itself.

**Third objection — epigenetic inheritance is an independent high-capacity channel.** *Transgenerational epigenetic inheritance demonstrates a heritable memory system independent of DNA.* Epigenetic inheritance is real, and the framework counts any transmitted organism-specific information it carries. The question is its demonstrated capacity and persistence. In mammals, extensive reprogramming occurs in the germline and early embryo, while known exceptions include imprinted loci, metastable epialleles, and RNA-mediated effects (Daxinger & Whitelaw, 2012). These establish additional bounded channels, not an unbounded second blueprint. A genome-scale independent program would predict stable, open-ended, DNA-invariant lineage divergence along its own informational axis; no natural epigenetic system has been shown to provide such organism-scale specification.

**Fourth objection — cortical inheritance and prions are DNA-independent blueprints.** *Ciliate cortical organization and prion conformations are inherited structurally and therefore refute a genome-centered specification budget.* These are genuine examples of non-DNA inheritance (Beisson & Sonneborn, 1965; Wickner, 1994). They demonstrate that realized structures can template continuation of particular structural or conformational states. Their known state spaces are bounded and low-dimensional relative to a full organismal microstate, however, and they do not provide an uncounted genome-scale trajectory program. They therefore illustrate state-carried inheritance rather than evade the framework: the inherited state is physically present, causally propagated, and counted as side information.

**Position within the formal apparatus.** No new machinery is required. Decompose $S_{\tau_0} = (Y_{\tau_0}, Z_{\tau_0})$ as in the scoped form of Lemma 1b. Fast residual degrees of freedom $Z$ mix, whereas slow organized variables $Y$ — organelles, spatial organization, marks, cortical structures, or conformational templates — are state-carried memory. Any organism-specific information in $Y$ is counted explicitly by Lemma 2 and by the zero-error side-information bound. The continuity of the cell lineage (*omnis cellula e cellula*) continues compiled state physically; it does not turn that state into an uncounted second program.

#### Backbone-only internal-coordinate calculation

For the 46-organism AlphaFold panel, the idealized backbone representation retains only the two torsions $\phi$ and $\psi$ per residue, while fixing bond lengths, bond angles, and peptide-bond geometry and omitting side chains and global pose. At angular resolution $\Delta\theta$, its direct structural cost is

$$B_{\text{torsion}}(\Delta\theta) = 2N_{\text{res}} \log_2\left(\frac{360^\circ}{\Delta\theta}\right).$$

The panel contains 241,282,135 residues and an aggregate coding input of 1.448 Gb.

Table S2: **Backbone-only internal-coordinate calculation across the 46-organism panel.** The representation retains only the two backbone torsions per residue and omits all side-chain and global-coordinate information.

| Angular resolution | Bits per residue | Aggregate backbone bits | Ratio to coding input |
|---|---|---|---|
| 45° | 6.00 | 1.448 Gb | 1.00 |
| 30° | 7.17 | 1.730 Gb | 1.19 |
| 20° | 8.34 | 2.012 Gb | 1.39 |
| 15° | 9.17 | 2.213 Gb | 1.53 |
| 10° | 10.34 | 2.495 Gb | 1.72 |
